# More accurate transcript assembly via parameter advising

**DOI:** 10.1101/342865

**Authors:** Dan DeBlasio, Kwanho Kim, Carl Kingsford

## Abstract

Computational tools used for genomic analyses are becoming more accurate but also increasingly sophisticated and complex. This introduces a new problem in that these pieces of software have a large number of tunable parameters which often have a large influence on the results that are reported. We quantify the impact of parameter choice on transcript assembly and take some first steps towards generating a truly automated genomic analysis pipeline by developing a method for automatically choosing input-specific parameter values for reference-based transcript assembly. By choosing parameter values for each input, the area under the receiver operator characteristic curve (AUC) when comparing assembled transcripts to a reference transcriptome is increased by 28.9% over using only the default parameter choices on 1595 RNA-Seq samples in the Sequence Read Archive. This approach is general, and when applied to `StringTie` it increases AUC by 13.1% on a set of 65 RNA-Seq experiments from ENCODE. Parameter advisors for both `Scallop` and `StringTie` are available on Github^1^.

## 1 Introduction

As the field of computational biology has matured, there has been a significant increase in the amount of data that needs to be processed and a corresponding increase in the reliance of users without computational expertise on the highly complicated programs that perform the analyses. At the same time, the number and sophistication of such tools has also increased. While the accuracy of such applications is constantly improving, a new problem has emerged: the sometimes overwhelming number of tunable parameters that each of these pieces of software brings with them. Changing an application’s parameter settings can have a large influence on the quality of the results produced (see Figure 1). When incorrect or non-ideal parameter choices are used, poorer results may be obtained or false conclusions may be reported.

**Figure 1:**
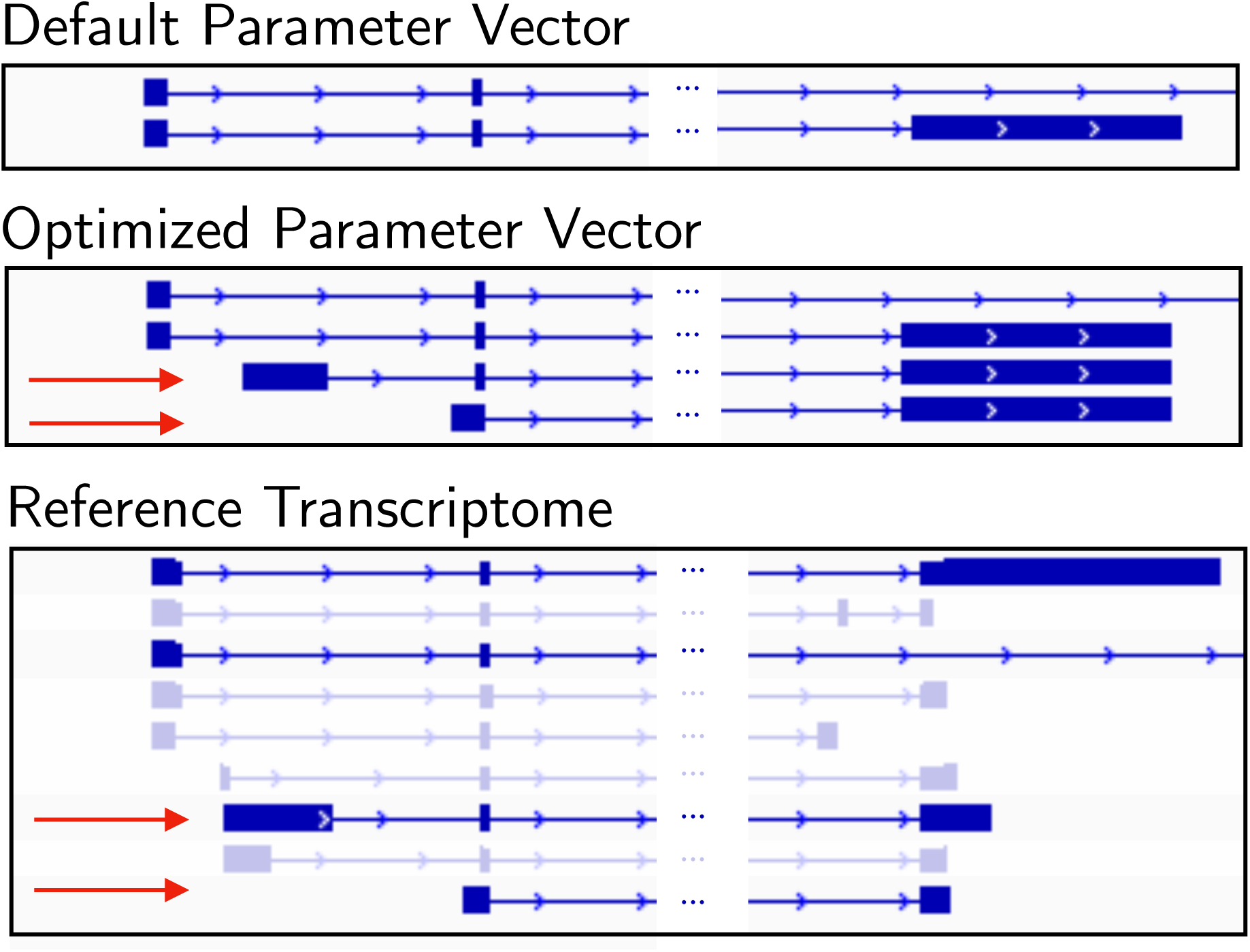
Influence of parameter choice on the produced transcriptome. The 3 sections of transcript assemblies are those assembled using `Scallop`’s default parameter vector, an optimized parameter vector, and the reference transcriptome for positions 30231125 to 30260786 on Chromosome 2 in `SRR543291`/`HISAT`. The red arrows highlight the two transcripts from the reference that are not recovered using the default parameter vector.

The default parameter choices that most users rely on for these programs are typically optimized by the algorithm designer to maximize performance on the average case. This can be a problem since the most interesting experiments are often not “average.”

Manually tuning the parameter settings of an application often produces more accurate results, but it is very time consuming. The tuning process can be accelerated for users with domain and/or algorithmic knowledge, as these experts can make more informed decisions about the correct direction to proceed when altering parameter values. But tuning the parameter choices to increase accuracy for one input does not imply that the results will be improved for all inputs. This means that, for optimum performance, tuning must be repeated for each new piece of data. In the case of high-throughput genomic analysis, this manual procedure is infeasible. For these applications, without some sort of automatic parameter choice system, the defaults must be used.

To address the automated parameter choice problem for multiple sequence alignment (MSA), **(author?)** [1] have defined a framework to automatically select the parameter values for an input. This process, called “parameter advising,” has been shown to greatly increase accuracy of MSA without sacrificing wall-clock running time in most cases, and it can readily be applied to new domains. Our new parameter advisor for transcript assembly depicted in Figure 2, details are provided in the next section.

**Figure 2:**
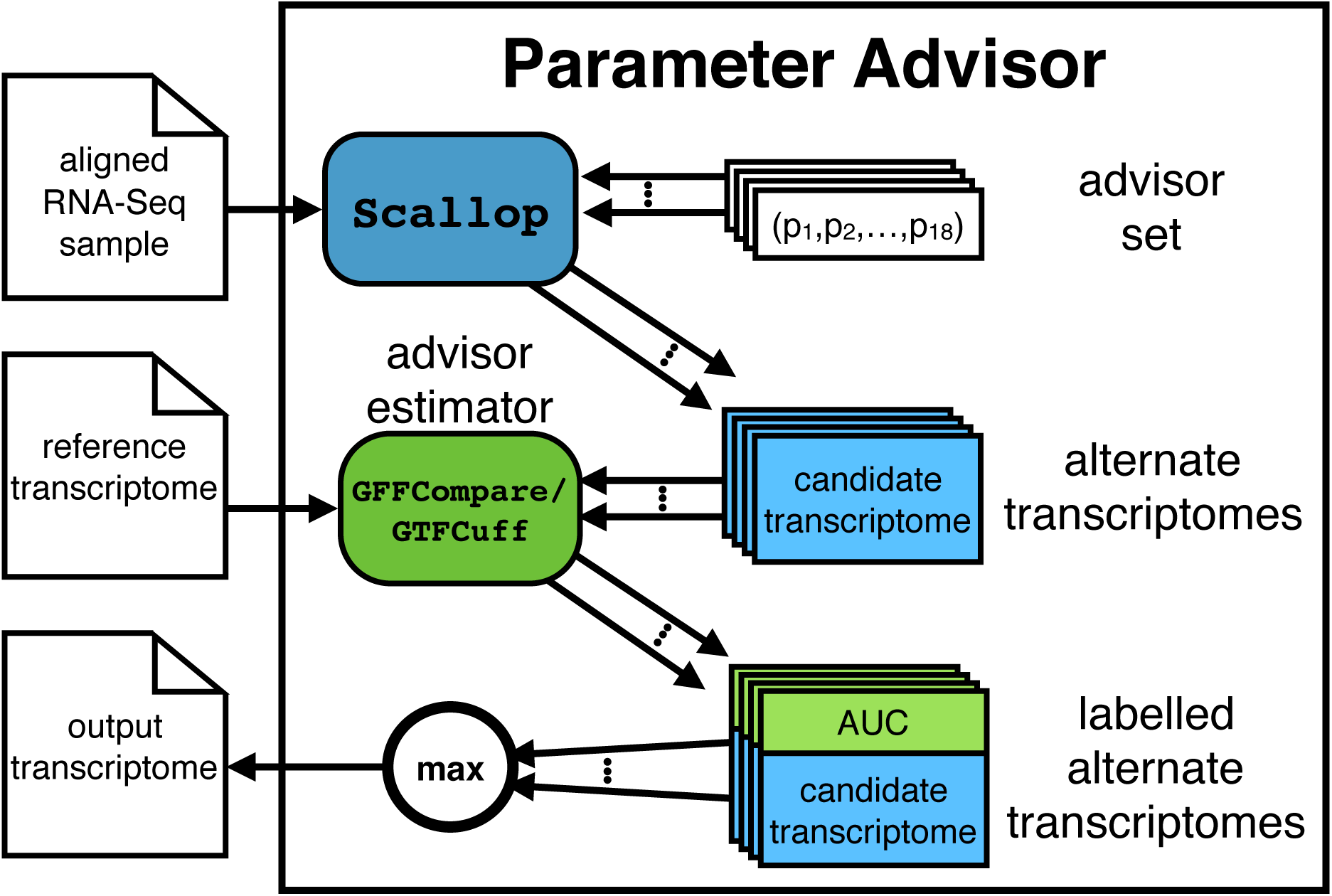
The Scallop parameter advisor. The advisor’s input mirrors that of `Scallop` (an RNA-Seq sample that has been aligned to the reference genome). A set of candidate assemblies is created by running `Scallop` on each parameter vector in the advisor set. The advisor then returns the assembly with the highest AUC value, obtained by running `GFFCompare` and `GTFCuff`.

In this work, we improve the quality of reference-based transcriptome assembly by extending parameter advising. Transcriptome assembly takes an RNA-Seq sample and a reference genome as input and reconstructs the set of transcripts that are present. Common tools for reference-based transcript assembly include `Cufflinks` [2], `StringTie` [3], `TranscComb` [4], and `Scallop` [5], Reference-based assemblers first align reads to the reference genome using a tool such as `HISAT` [6], `STAR` [7], `TopHat` [8], or `SpliceMap` [9]. Using the read splice locations (the positions where a read maps to non-neighboring locations on a genome), the assembler constructs the exons and splice-junctions of each transcript. The produced transcriptome consists of a combination of transcripts that can be mapped to ones we already know and transcripts that are unique to the sample that was assembled. These transcriptomes are used to perform analyses such as expression quantification [10, 11] and differential expression [12, 13].

For transcriptome assembly, the quality of a program’s output is commonly measured using the area under the receiver operator characteristic curve (AUC) when the produced transcripts are mapped to a reference transcriptome. We will use this as our method for selecting parameter choices for a given input.

### Contributions

The contributions of this work are threefold: first, we show for the first time that advising sets of parameter vectors can be constructed for tools with large numbers of tunable parameters; second, we take some of the first steps towards producing a fully automated transcript analysis pipeline by automating sample-specific parameter selection for multiple applications; and third, we show that AUC is a better measure to use for parameter optimization in reference-based transcript assembly than existing other *de novo* metrics.

We show that by applying the parameter advising framework, we can greatly increase the quality of the transcriptomes produced using the `Scallop` assembly tool. Using this approach, the area under the curve shows a median increased of 8.7% over using only the default parameter vector on a set of 10 RNA-Seq experiments contained in the ENCODE database that are commonly used for benchmarking. In a high-throughput pipeline the median improvement is even larger, 28.9% higher AUC than the default parameter vector on over 1500 samples from the Sequence Read Archive.

We also confirm that this method can increase AUC for other programs by applying it to `StringTie`, another popular reference-based transcript assembler. For a set of 65 examples from the ENCODE database, we are able to increase its AUC by 13.1% over using only the default parameter vectors.

## 2 Developing a parameter advisor for transcript assembly

A parameter advisor has two components: (1) a set of parameter vectors – assignments of a value to each of the tunable parameters for the application – called an “advisor set”; and (2) an assessment criteria – a method to rank the quality of multiple solutions – called an “advisor estimator”. The advisor selects the appropriate parameter vector by first running the application on the input using each parameter vector in the set, and selecting the parameter vector that produces the best result according to the accuracy estimator. The instantiations of the application being tuned are independent processes that can be executed in parallel. Assuming that the number of processors available is at least the number of parameter vectors in the advisor set, the only additional wall time is the assessment of the results using the accuracy estimator (which can also be performed in parallel) and the comparison of these values, both of which are negligible compared to the running time of the application in most cases.

Parameter advising is an example of *a posteriori* parameter selection — it examines an application’s output to select a parameter setting. In contrast, separate work has been done in other fields on *a priori* selection, where the parameters are chosen in advance by looking at the raw input alone, or a subsample of the input along with its result. This includes work such as `SATZilla` [14] for choosing from a collection of SAT solvers, `ParamILS` [15] which finds optimal settings for the `CPLEX` computational optimization tool, `KmerGenie` [16] for finding appropriate *k*-mer sizes for genomic assembly, as well as many tools developed for tuning hyperparameters in machine learning such as `TPOT` [17] which uses genetic algorithms and `Spearmint` [18] which uses bayesian optimization. *A priori* prediction is necessary in cases when it is not feasible to apply multiple configurations, but more information is available when performing *a posteriori* assessment since the full final solution can be examined.

### Advisor estimator

For transcriptome assembly, a natural choice for the estimator is the area under the curve (AUC) of an assemblies prediction of known transcripts. The minimum transcript coverage value is used to rank the transcripts, then label the set of assembled transcripts that match with the reference as positives and all others as negatives. The minimum coverage of a transcript is the minimum number of reads that are aligned to each position along the transcript’s length.

In the case of transcript assembly, the ground truth set (the reference transcriptome) is much larger than we will ever see in any single sample since it contains all transcripts that have been identified and verified. Therefore, the sensitivity of any transcript assembly will by definition be very low, in most cases below 0.1%. This means that the AUC value will also be very low. But, since there are close to 200,000 transcripts in the reference, even small differences in sensitivity are large in the true number of transcripts recovered correctly.

### Advisor set

Finding an advisor set is especially challenging because `Scallop` has 18 tunable parameters compared to approximately 5 for multiple sequence alignment, the previous application of parameter advising. This means the previously developed method of enumerating a parameter vector universe and then using combinatorial optimization to find an advisor set is infeasible. The `Scallop` transcript assembler generates a transcriptome from a set of reads that have been aligned to a reference genome. It first splits the genome into regions of non-overlapping reads, which are called bundles. These bundles can be thought of as genes or groups of overlapping genes. Then, within each bundle a splice graph is constructed based on the split reads that define possible exon boundaries. Paths through the splice graph define potential transcripts, and the final set of transcripts is formed by decomposing the splice graphs into paths while trying to respect as many of the read mappings as possible. The tunable parameters of `Scallop` govern various stages of this process, but we treat the application as a black box and do not examine the actual function of each parameter only how the manipulation of it’s value impacts transcript quality.

### 2.1 Analyzing parameter behavior

We calculate the AUC of a transcript assembly by allowing `Scallop` to output all predicted transcripts rather than thresholding on on minimum transcript coverage (the average number of reads that are aligned to each position along the transcript’s length), and then allowing the tools GFFCompare^2^ and GTFCuff^3^ to calculate an AUC by thresholding the minimum transcript coverage value. Because the reference transcriptome contains a large number of very rare transcripts the sensitivity is very low, and in turn so is AUC. Since we may not know ahead of time for any given input which transcripts, or even how many, to expect in a sample we cannot use a reduced reference to inflate the sensitivity. While the values are small, comparing them will still indicate relative performance improvements with respect to number of true and false positives using various parameter choices. In this work, as opposed to **(author?)** [5], we choose to not separately evaluate multi-exon transcripts and single-exon transcripts but rather maximize a combined AUC.

Iterative optimization strategies such as gradient ascent [19], simulated annealing [20], and coordinate ascent [21, sec. 5.4.3], work by systematically searching high-dimensional spaces based on a specific optimization criteria. For these methods to work well the parameter landscape should be free from a large number of local maxima as well as large discontinuities.

To determine the relationship between AUC and the value of the `Scallop` parameters, we calculate the AUC of the assemblies produced when varying a single parameter’s value while keeping the remaining parameters at the default. Figure 3a shows the effect of varying the “minimum subregion gap” and “minimum transcript length, base” parameters. Figure 3b shows the relationship between the parameters and AUC when varying both at the same time. Note that throughout this work, we multiply area under the curve values by 10^4^ for ease of presentation. AUC is a value in the range [0, 1], but generally for transcript assembly the value is very small, typically < 0.1.

**Figure 3:**
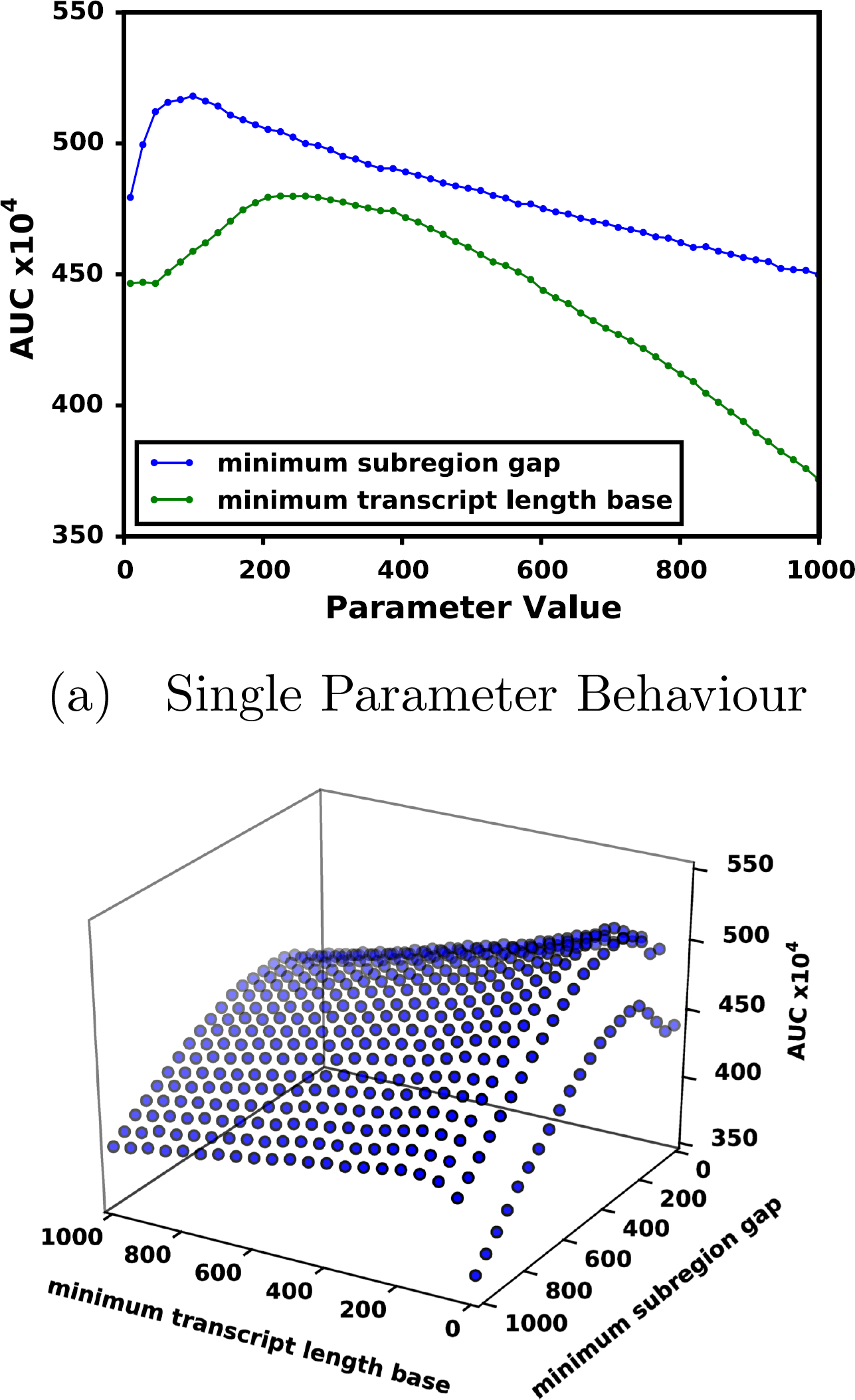
AUC for various values of the \minimum subregion gap” and \minimum transcript length base” parameters. The points in the plot show the area under the curve (vertical axis) for the transcriptome produced by changing the value each of the parameters either (a) alone or (b) together leaving all other parameters their default values on

We examined the parameter behavior curves for several experiments from the ENCODE database, and found that the curves for all 16 continuous parameters and all of the pairs of parameters tested contained only one visible local maximum. These tests suggest that there may be very few local maxima in the high-dimensional parameter space which means iterative optimization procedures are less likely to get stuck at poor local maxima.

### 2.2 Finding an advisor set using coordinate ascent

The greedy coordinate-ascent-based procedure that we use here starts at the default parameter vector. One dimension (parameter) at a time, we examine the AUC of the parameter vector with that parameter changed by one step in each direction and update our current vector if we see an improvement. We continue tuning one dimension until no more improvements are made. Our procedure is deterministic meaning we would never take a step that decreases (or maintains) AUC, unlike many implementations of coordinate ascent which include some randomness in decision making. The step size for each dimension has a time-versus-granularity tradeoff: the larger the step size, the less time spent in low AUC regions of the landscape; but when the size is too large, the maximum may be repeatedly stepped over without ever being found. Rather than use a computed step size for every coordinate in each iteration like many implementations of coordinate ascent, we use predetermined but decreasing step sizes, taking inspiration from simulated annealing. We start with different step sizes in each dimension that are large relative to the default value for that parameter, and any time we interrogate the whole set of parameters without making any change we decrease all of the step sizes by a factor of 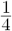 and repeat the process. We continue optimizing the parameters in this manner until both: (1) all of the step sizes are small (1 for integer parameters and 0.01 for real numbers) and (2) no more improvements can be made within one step from the current parameter vector. For the tunable parameters in `Scallop` that accept only binary input, we tested both options each time the parameter was explored.

Coordinate ascent will find parameter vectors with higher AUC for an input, but it is slow so it is not a viable procedure for finding input-specific parameter choices in practice. Instead, we can use coordinate ascent to find higher AUC parameter vectors for a set of examples, then use the collection of optimal parameters as the advisor set. Since the advisor sets are computed in advance so as long as the the set of examples we use is diverse, meaning it represents the range of possible inputs, the advisor sets can be reused for any new input.

### 2.3 Data

We use 3 sets of human RNA-Seq experiments to train and validate parameter advising:

- `ENCODE10` contains a collection of 10 RNA-Seq experiments from the ENCODE database [22] that were used to benchmark `Scallop` and have been extensively used to evaluate transcriptome assembly tools [3, 4]. 30 examples were produced by aligning each sample to the human reference genome (GRCh38) using three tools: `HISAT`, `STAR`, and `TopHat`. A subset of these examples was used to find the `Scallop` default parameter vector.
- `ENCODE65` contains a collection of 65 RNA-Seq experiments, also from ENCODE, that were not included in `ENCODE10` and that had preexisting alignments in the database. These alignments are produced using an aligner selected by the group that submitted the sample and are mapped to either GRCh37 or GRCh38.
- `SRA` contains a collection of 1595 RNA-Seq experiments from the Sequence Read Archive [23] that have been filtered for quality. We eliminate any sample that contained very few reads unaligned (< 1GB sequence file) or aligned (< 1GB alignment file) since this is an indication that the experiment may be degraded in some way. All remaining samples were aligned using `STAR` to GRCh38.

All of experimental identifiers and command line arguments are available at: https://github.com/Kingsford-Group/scallopadvising

## 3 Validating the transcript assembly parameter advisor

### 3.1 Finding a Scallop advisor set

The `ENCODE10` dataset is reasonable for training because it is highly diverse and is expected produce parameter vectors that should generalize. It contains samples that are widely accepted as benchmarks and has examples that have been generated using a collection of commonly used aligners. The coordinate ascent procedure described in Section 2.2 was used to find improved parameters for each sample. While all parameter were tuned when optimizing parameter choices, for some parameters the final parameter vectors never included non-default parameter values, namely “maximum dynamic programming table size”, “maximum edit distance”, and “minimum router count.”

Most of the parameters values deviate quite far from each other between samples, meaning there is unlikely to be one parameter choice that works well for all of the training examples. Given the output of the optimization, a new default parameter setting could be recommended for some single parameters, such as in the case of “minimum mapping quality” where almost all samples used a value of 11 rather then the default of 1, and “minimum transcript length increase” where most samples found improvement by selecting values that are much smaller than the default. In fact, we find that the parameter vector found for SRR545723/`TopHat` had higher AUC for all of the samples in the `ENCODE10` and would be the best vector to use as the default, 533.2 on average versus 475.5. The deviation of the parameter vectors from the default is not surprising given that in this work we are optimizing AUC on all transcripts, rather than only multi-exon transcripts as was done previously. When training we placed no restrictions on any of the ranges of values any particular parameter can take on, so while the values may seem somewhat unintuitive to domain experts they are the values that give the highest AUC value for the training example.

In reduced resource environments (e.g., when 31 processors are not available), it may be desirable to run fewer parameter settings to keep the number of parallel processes smaller than the number of available threads. We used the oracle set-finding method described by **(author?)** [1, 24] to find a subset of parameter vectors that maximizes the average AUC for advising.

Advisor subsets are found using an integer linear program that has two sets of binary variables: one variable for each parameter vector, and one for each example/parameter pair. Where an example/parameter pair is a parameter vector used to assemble an example. Constraints are used to ensure that only one pair for each example is chosen and that the associated parameter setting is also chosen. The objective is then to maximize the sum of the accuracies of the chosen pairs while only selecting a predefined number of parameter vectors. Using the samples in `ENCODE10`, we found advising subsets of 1, 2, 4, and 8 parameter vectors. The advising subset choices are shown Table 1. Note that an advising set of size 1 is equivalent to finding a new default parameter vector since it maximizes the average accuracy across the training examples.

**Table 1:**
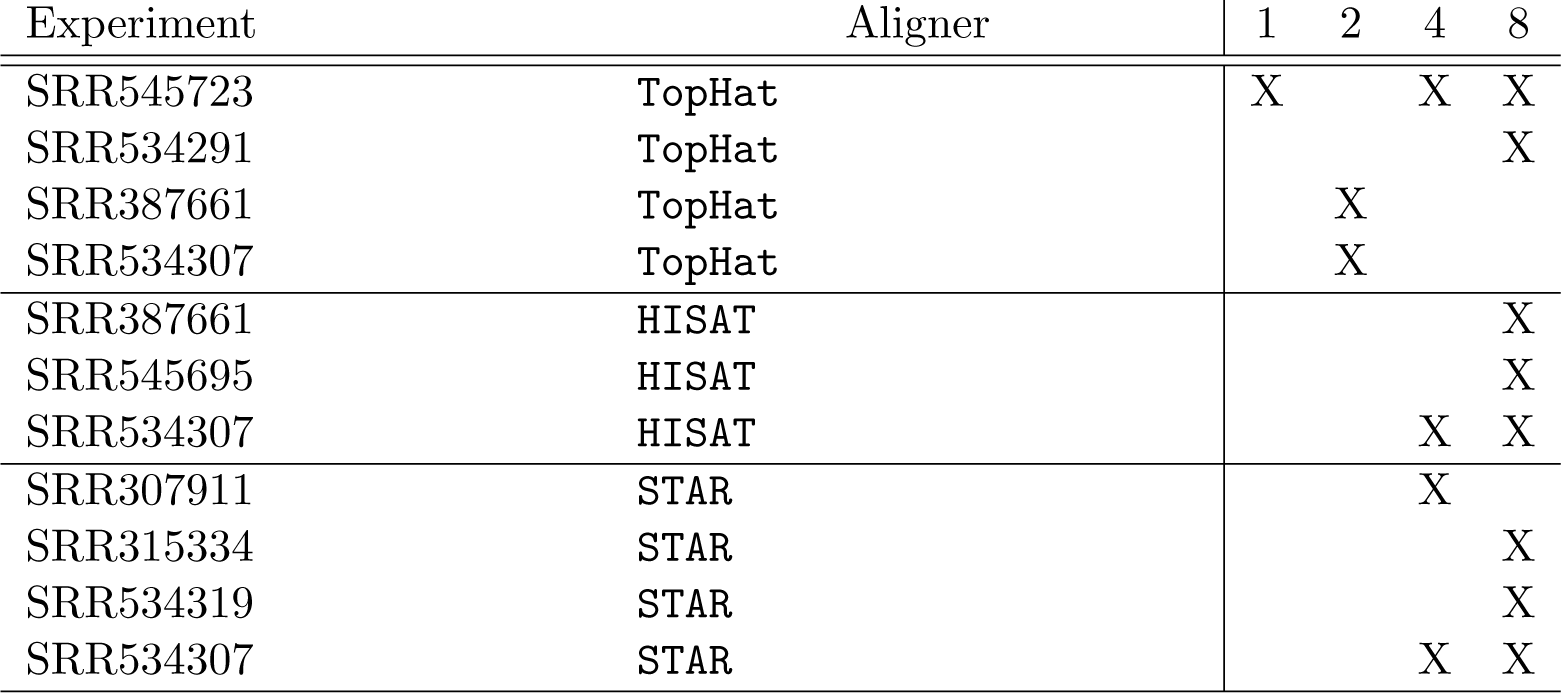
Parameter vector subsets

#### 3.1.1 Advising on the training set

Figure 4 shows the AUC for all of the samples from `ENCODE10`. For each example (a sample combined with an aligner) there are two values shown: the AUC of the parameter vector produced as a result of coordinate ascent, and the AUC of the leave-one-out advising parameter vector. For the leave-one-out experiment, advising was limited to the 18 parameter vectors that were learned on examples produced using the 2 aligners and 9 samples that were different from the example being tested. This test shows that the parameter vectors learned on specific examples, can generalize and improve the AUC for other unrelated examples.

**Figure 4:**
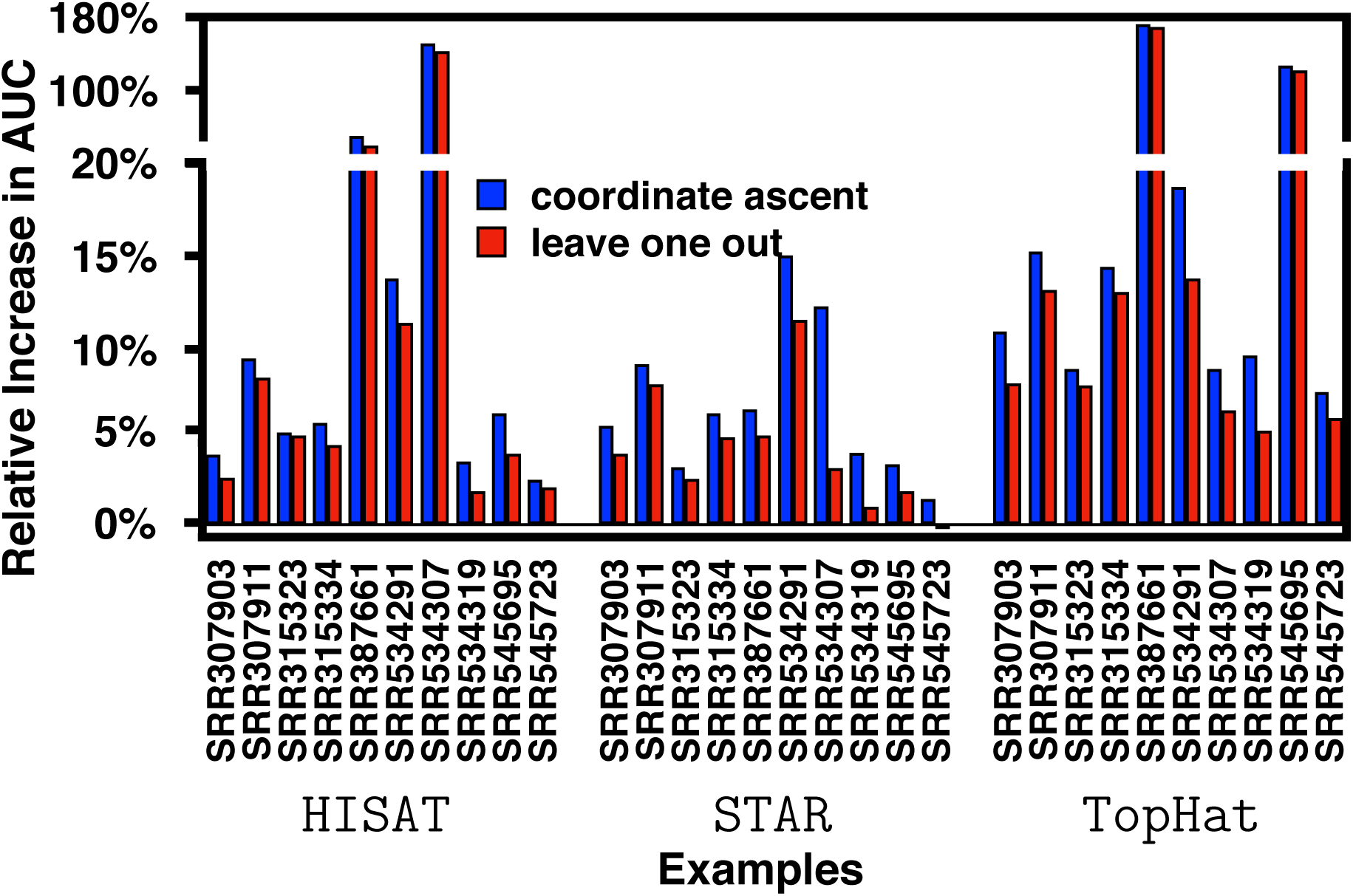
Increase in AUC for examples in ENCODE10. Each of the 30 examples is listed on the horizontal axis. Each bar’s height is the normalized difference between the default AUC and that of the assembly produced using either coordinate ascent (blue) or leave-one-out (red).

### 3.2 Assessing the generality of learned parameter vectors

#### 3.2.1 ENCODE65

The `ENCODE65` dataset is used to show that on a large number of samples from a range of aligners (possibly using non-default parameter settings) advising for `Scallop` provides a higher AUC transcriptome. Figure 5 shows the relative increase in AUC, the AUC of the transcript assembly produced by the advised parameter vector normalized by that of the default parameters, for all 65 examples in `ENCODE65`. Using advising on this highly diverse set of samples increases the AUC of each transcriptome by a median of 31.2%. When the default parameter vector performs well there is a smaller increase in AUC. This is reasonable since there is less room for improvement for these samples. When using the resource limited sets, the median increase remains 18.2%, 19.0% and 24.4% for sets of 2, 4, and 8 parameter settings respectively. Even with these small sets, there is a large increase in AUC.

**Figure 5:**
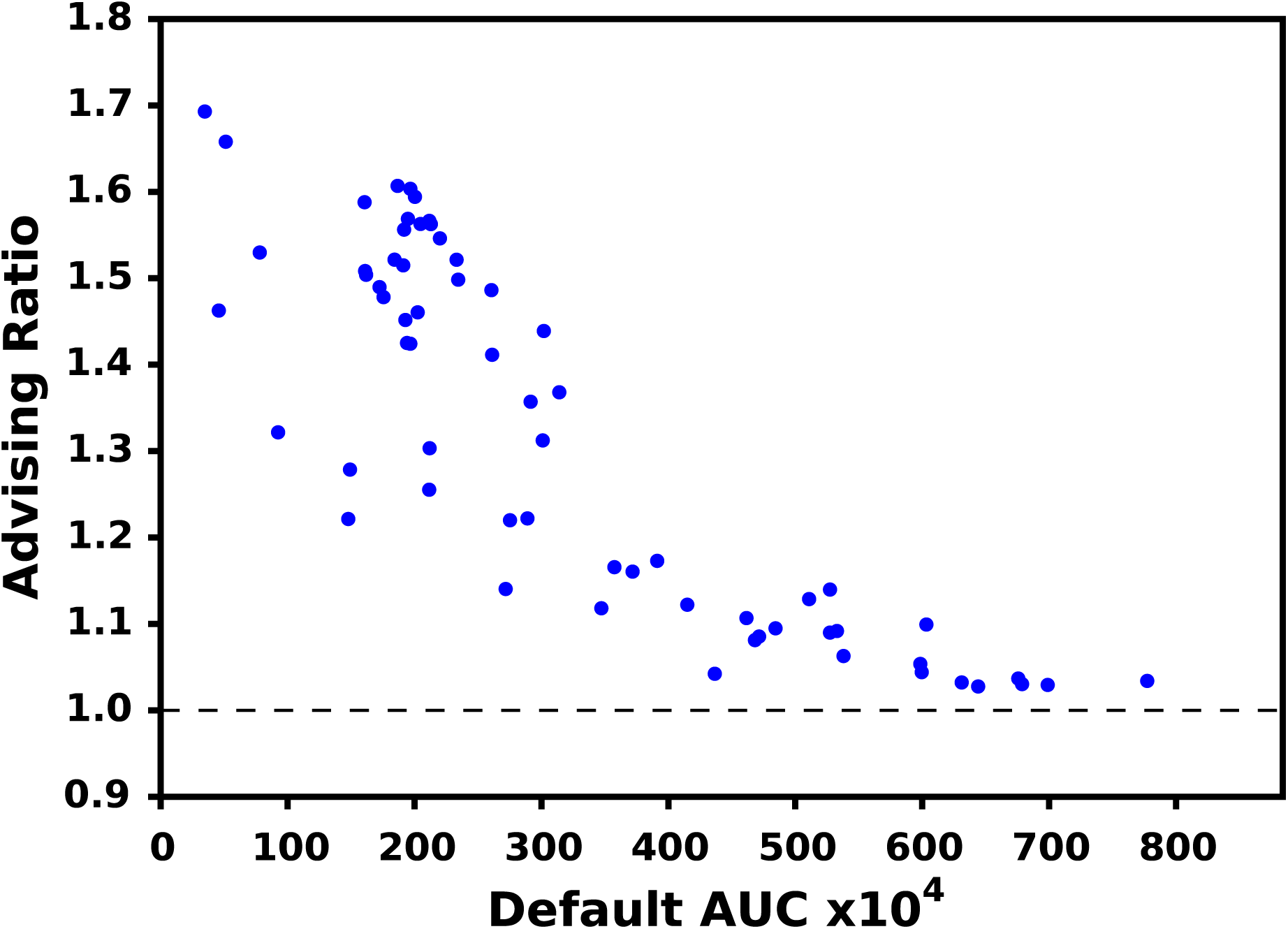
AUC improvement for ENCODE65. Each point is a single experiment positioned by the AUC of the default `Scallop` parameters (horizontal axis) and the ratio of the advised AUC over the default (vertical). A value above 1.0 indicates an improvement.

Figure 6 shows the frequency with which each of the 31 (the whole set of parameter vectors learned on `ENCODE10`), 8, 4, and 2 parameter vectors provides the maximum AUC when running parameter advising on the `ENCODE65` set. Not all 31 parameter vectors provide the best AUC over the 65 samples, but more parameter settings are maximal across the samples as you increase the size of the advisor set. Only 4 parameter vectors of the set of 8 provide an assembly with the maximum AUC for all of the samples in `ENCODE65`, as well as only 2 of the 4.

**Figure 6:**
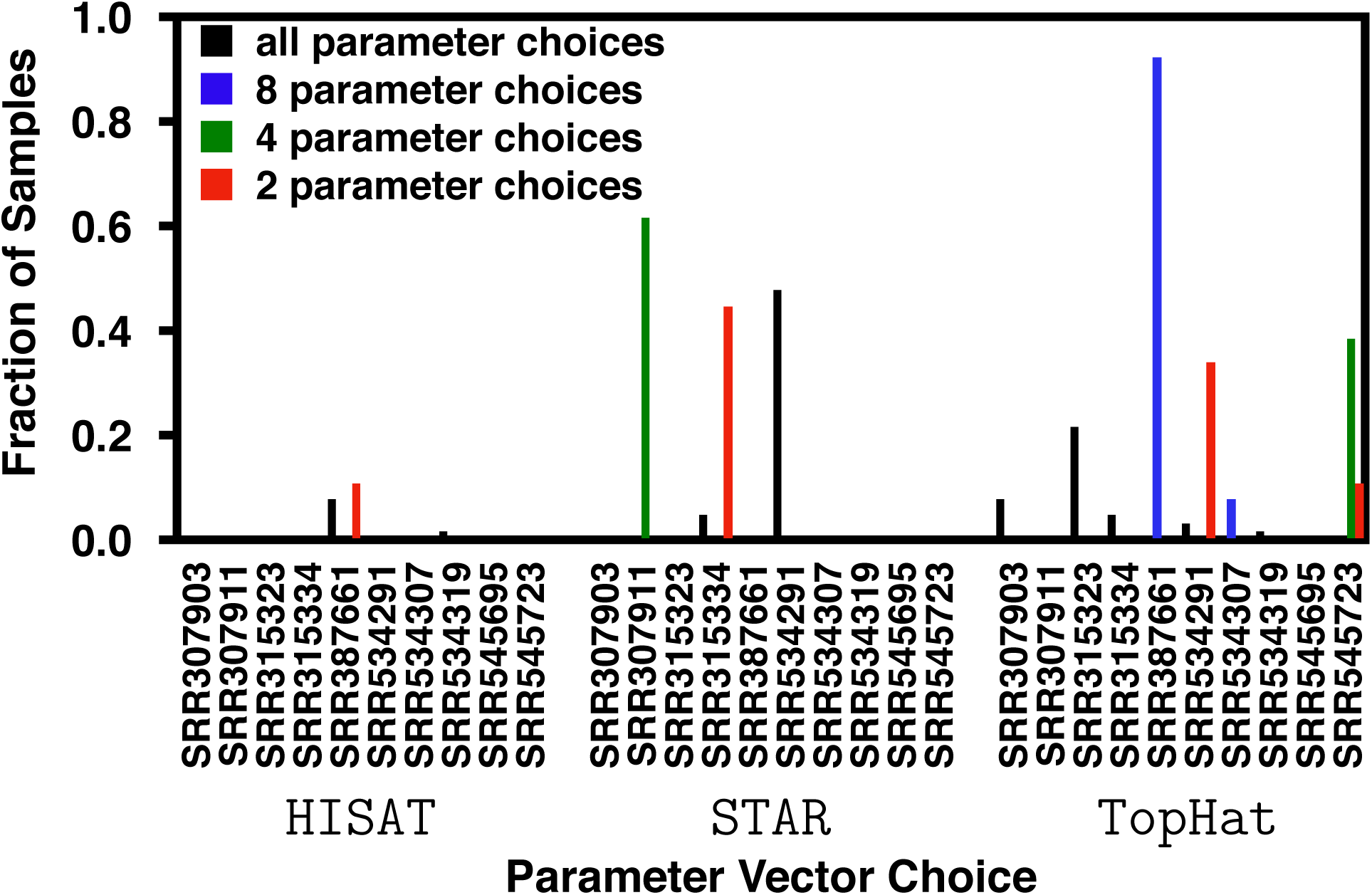
Frequency of parameter vector use for advising the samples of ENCODE65. The horizontal axis shows labels of the experiments from which the parameter vectors were trained that produced the highest AUC transcriptome for any sample in the dataset. The vertical axis is the fraction of samples that have that parameter vector as the maximum. The four groups of bars show the use in the full set of 30 parameter vectors and the reduced sets of 2, 4, and 8 parameter vectors.

##### Random Advisor Sets

To confirm that the increase in accuracy is due to our advisor set construction method and not an artifact of having multiple choices of parameter vectors, a collection of random parameter vectors were generated and used for parameter advising. A range was defined for each tunable parameter by examining all of the values that provided an increase in AUC for any example at any stage in coordinate ascent. A random vector was then constructed by selecting parameter values for each parameter uniformly at random in these ranges. In total, 30 such parameter vectors were generated to match the advisor set size developed using coordinate ascent. This randomization procedure was then replicated 100 times to ensure stability of the average.

Figure 7 shows the AUC achieved by parameter advising on `Scallop` using the the coordinateascent-derived advising set versus the AUC of advising using the random advisor sets. The default parameter vector was left out of the all advising sets. Because of this, many of the randomly generated advisor sets (29 of 65) led to a decrease in accuracy relative to the default. On some examples the performance is similar between the two sets, but the average increase in AUC is much higher for the coordinate ascent advisor set (median AUC increase of 31.23 versus 5.59). In all of the 65 examples, the coordinate ascent sets outperform the random ones.

**Figure 7:**
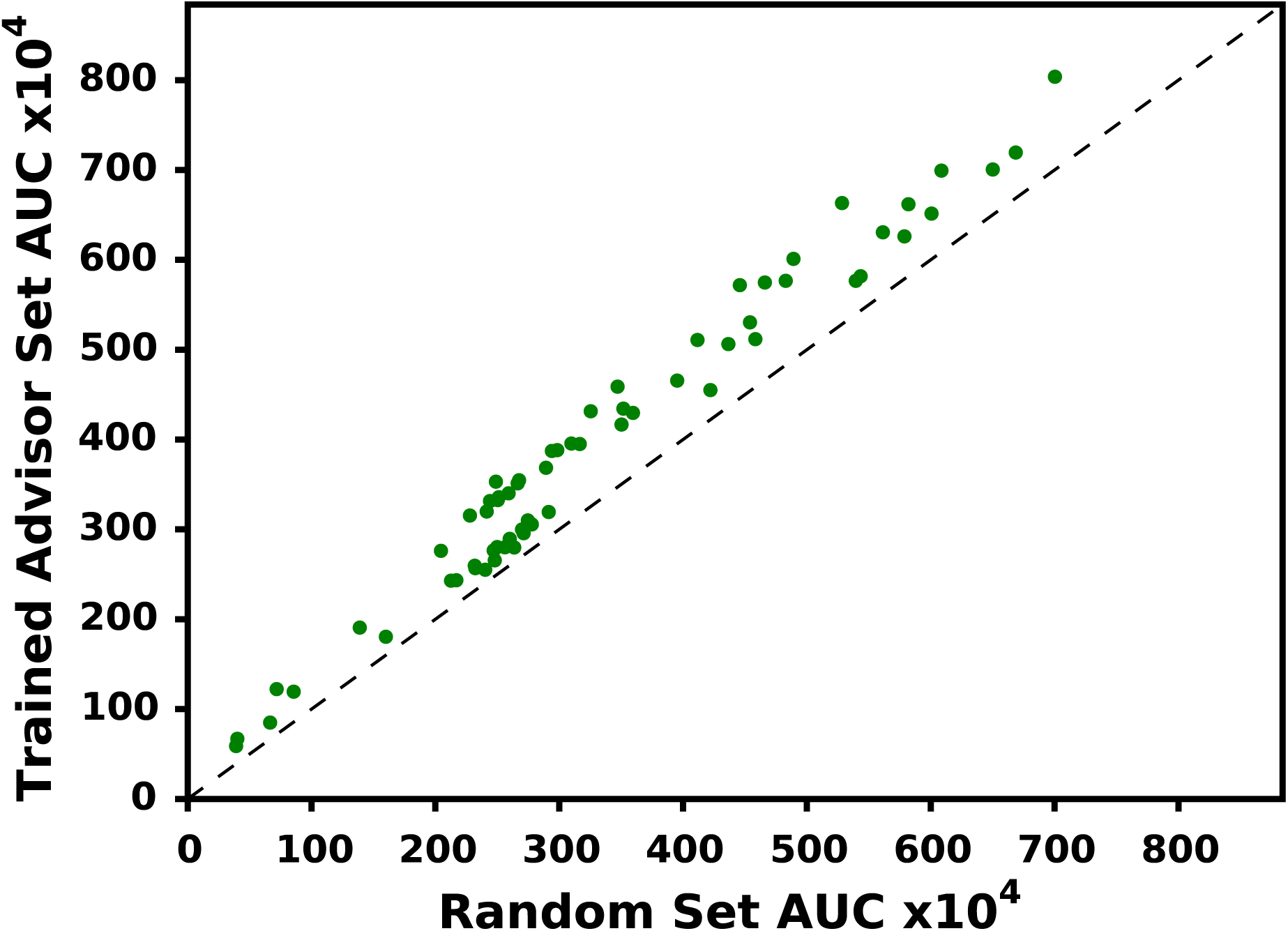
Comparison to randomly generated advisor sets. Each point in the chart represents one example from `ENCODE65` positioned by the AUC value of using a randomly generated advisor set (horizontal axis) or the trained advisor set (vertical axis) averaged over 100 random replications. Points above the diagonal line show an improvement of training over random.

#### 3.2.2 SRA

Our `SRA` dataset gives some insight into the improvement that can be gained in a high-throughput environment. It contains a large number of samples that have all been preprocessed in the same way with respect to the aligner. The advising ratio for the 1595 samples in `SRA` is shown in Figure 8 compared with the AUC for the transcriptome produced using the `Scallop` default parameter settings. Because, in general, the samples in `SRA` have a smaller initial area under the curve than those in Figure 5 (AUC values of 241.3 and 325.0 respectively), the median improvement is higher at 28.9%. For several samples in the set the AUC increases by more than a factor of 3. These improvements are also seen for the resource-limited advisor sets where the median improvement is 25.6%, 24.1% and 24.3% with 2, 4, and 8 parameter vectors, respectively. Notice that the increase in AUC actually goes down slightly when increasing the size from 2 to 4. This is likely an artifact of the reduced advisor sets not being subsets of each other. This means that the parameter vectors and sets may be slightly overfit to the training data. Figure 9 shows the frequency with which each of the 31, 8, 4, and 2 parameter vectors provides the maximum AUC when running parameter advising on the `SRA` set. Because all of the examples in `SRA` were aligned using the same aligner many of the choices are maximal more frequently. Surprisingly, even though all of the examples are aligned using `STAR`, many of the higher-frequency parameter vectors had been optimized for examples that were aligned using `TopHat` (IDs 1–10). Ties, if any existed, would be resolved in alphabetical order of the experiment name then aligner but no ties for the maximum AUC were found in `SRA` or `ENCODE65`.

**Figure 8:**
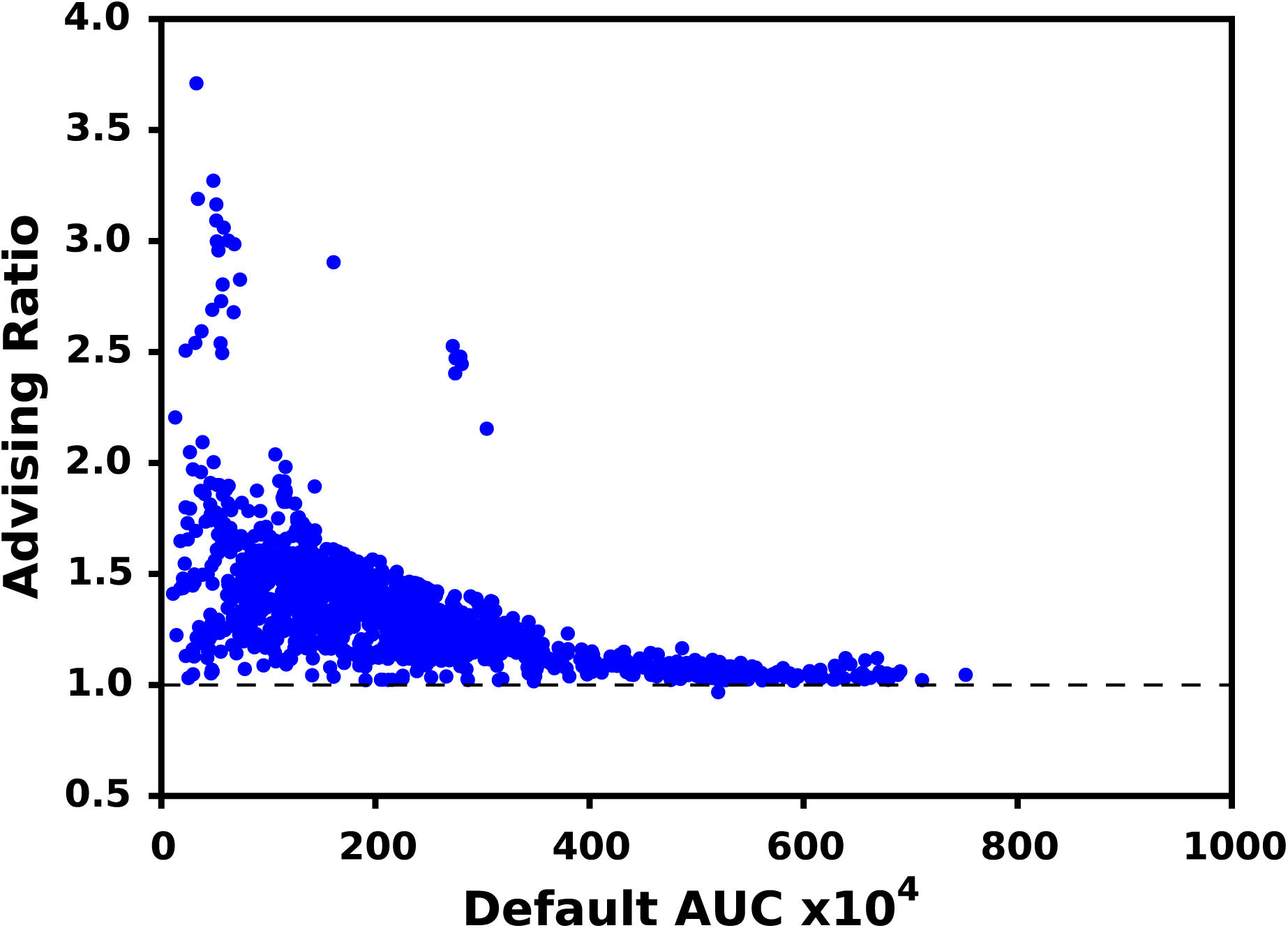
AUC improvement the SRA assemblies. Each point is a single experiment positioned by the AUC of the default `Scallop` parameters (horizontal axis) and the ratio of the advised AUC over the default (vertical). A value above 1.0 indicates an improvement. For this test the default was excluded from the advising set, but it can be included in practice to ensure the AUC is never reduced.

**Figure 9:**
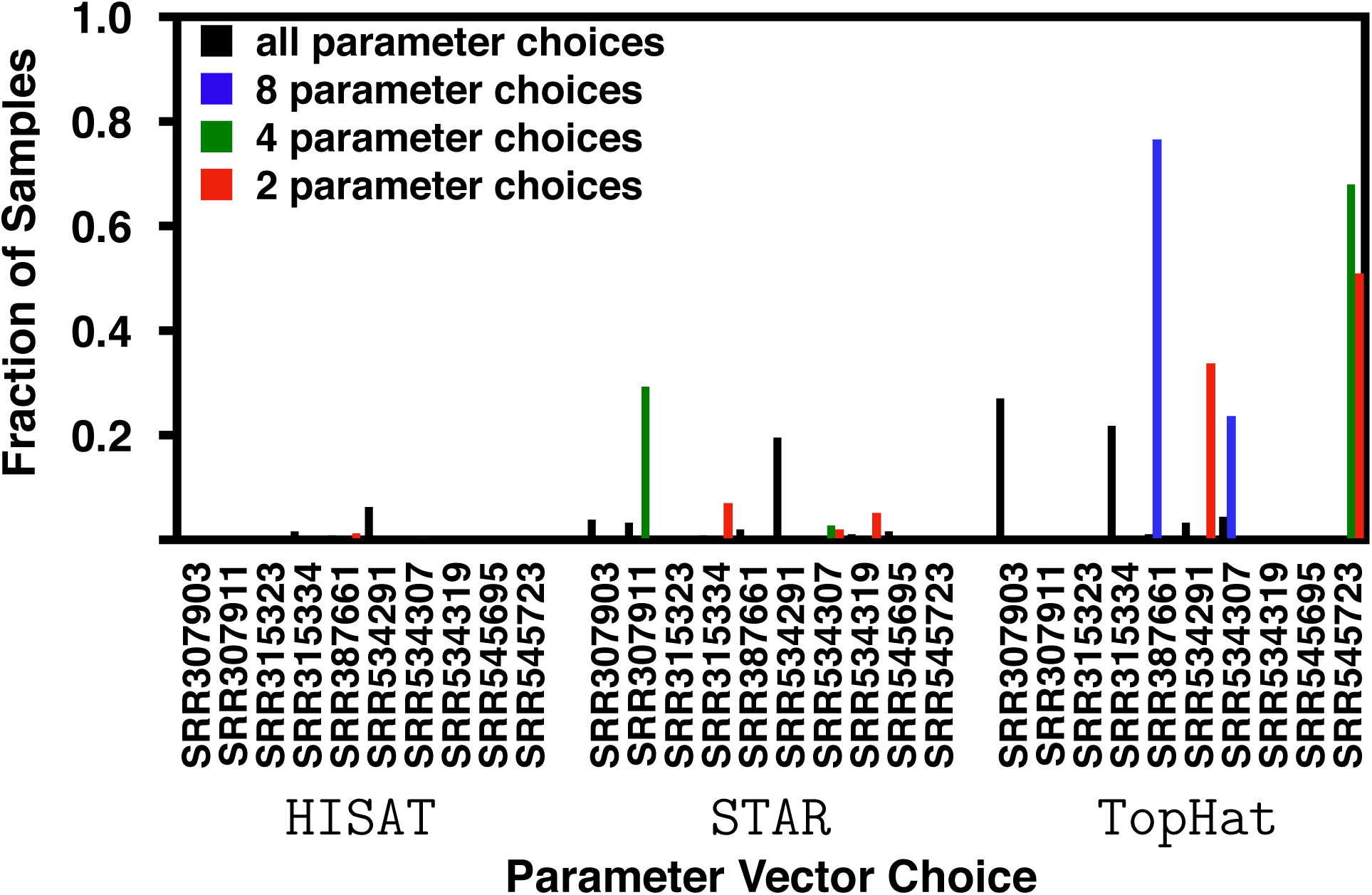
Frequency of parameter vector use for advising the samples of SRA. The horizontal axis shows labels of the experiments from which the parameter vectors were trained that produced the highest AUC transcriptome for any sample in the dataset. The vertical axis is the fraction of samples that have that parameter vector as the maximum. The four groups of bars show the use in the full set of 30 parameter vectors and the reduced sets of 2, 4, and 8 parameter vectors.

### 3.3 Running time

The wall time of running coordinate ascent is much larger than the running time of any single instance of `Scallop`. For the 30 examples from `ENCODE10`, running coordinate ascent for any single example took between about 40 hours and over 22 days.

Running an advisor would take only as much wall-time as running a single instance of `Scallop` with appropriate available resources. Running `Scallop` using the default parameter vector for these same samples takes between ∼7 minutes and 1 hour. Even if no parallelization was possible, parameter advising would be able to run in a fraction of the time of running coordinate ascent.

### 3.4 Advising for StringTie

In order to show the generalizability of this method, we also applied it to the `StringTie` transcript assembler. As before, we ran coordinate ascent on the 10 experiments in `ENCODE10`, now using `StringTie`, to select the 30 non-default parameter vectors. Since `StringTie` has only 9 tunable parameters, the coordinate ascent time was much shorter, but the increase in accuracy was still observed. For the 30 coordinate ascent runs, we saw a median increase in AUC of 12.2% on `ENCODE10`. We also saw 10.0% increase in AUC on the similar leave-one-out experiments as those performed above.

Figure 10 shows the advising ratio for the 65 RNA-Seq samples from `ENCODE65`. For these examples the median gain in AUC is 13.1% over using only the default parameter vectors. For the `StringTie` assembler, samples with lower AUC using the default parameter vectors still generally have higher advising ratios but this correlation is not as strong as with `Scallop`.

**Figure 10:**
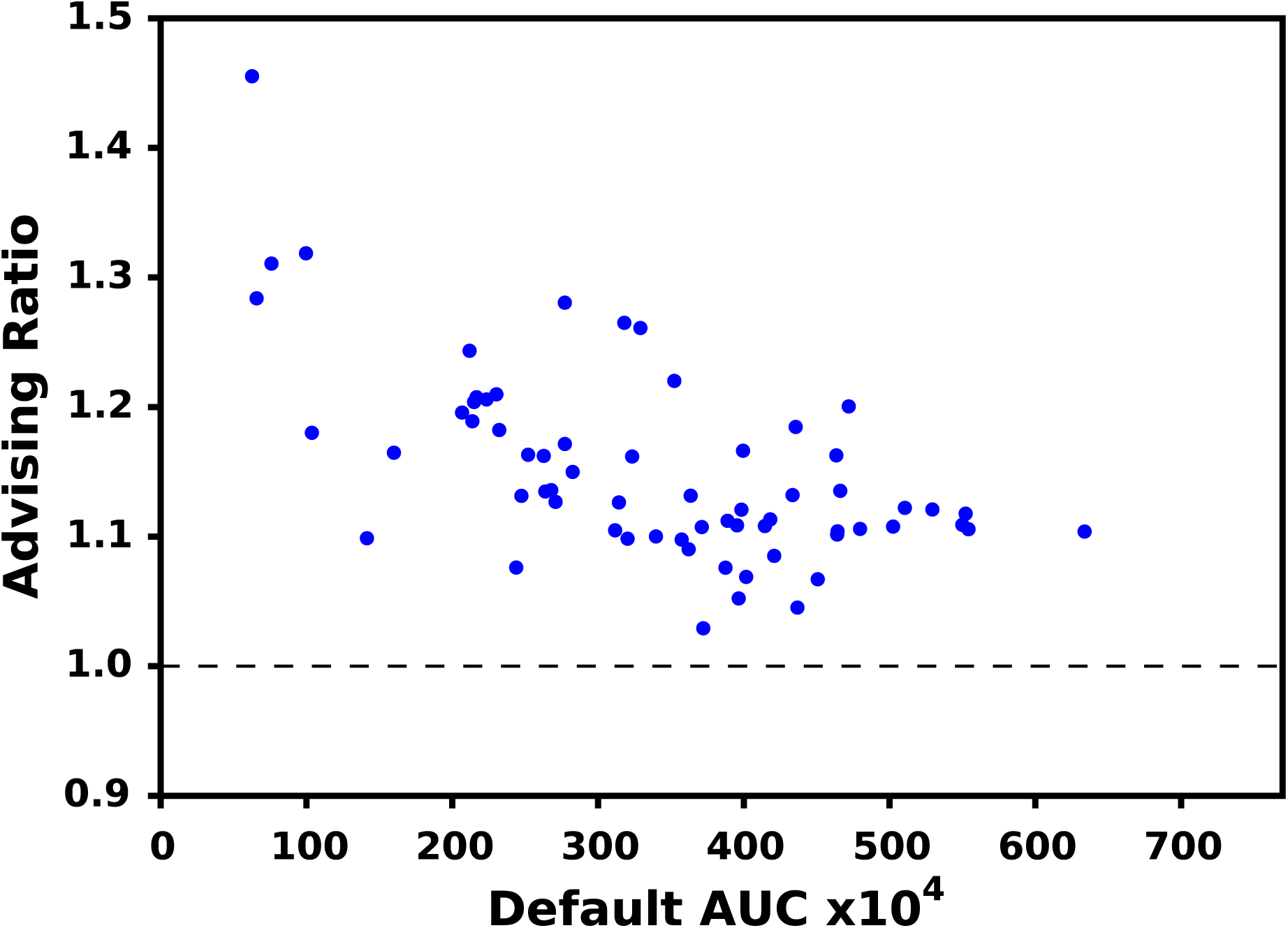
AUC improvement the ENCODE65 assemblies using StringTie. Each point is a single experiment positioned by the AUC of the default `Scallop` parameters (horizontal axis) and the ratio of the advised AUC over the default (vertical). A value above 1.0 indicates an improvement.

### 3.5 Justification for using AUC as the advising accuracy estimator in parameter optimization

Comparison to the reference transcriptome is the commonly used metric to benchmark reference-based assemblers. However, the performance of AUC is bounded by the completeness of the reference transcriptome. By definition, all transcripts in an assembly that do not map to the reference provide a reduction of AUC, even those that represent “novel” transcripts. Here a novel transcript is one that is present in the sample but has not been included in the reference yet.

We tested the ability to improve transcript assembly parameter choices of 3 alternatives to AUC which do not rely on the reference transcriptome in simulation where we know the total ground truth. The metrics we compare to are `Transrate` [25], “number of reads” mapped to the transcriptome using `Salmon`, and a linear combination of the features from `Transrate`, number of reads, and other novel features combined as a weighted sum (labeled as “linear”).

Simulated datasets were constructed using the samples in `ENCODE10`. We restricted the reference transcriptome for each experiment to be the collection of transcripts from the reference that map to assembled transcripts in that experiment. We then limit the set of sequencing reads to those that map to the transcripts in this reduced reference.

Since the reduced reference has all of the transcripts in the sample and nothing else, the AUC using this reduced reference transcriptome in this case is the ground truth accuracy. We can then compare just how well the other metrics are able to recover this known truth set.

A subset of this reduced reference set was used to test AUC in coordinate ascent. In this way, AUC only has partial information about which transcripts are in the sample.

Figure 11 shows the improvement of the parameter vectors found using coordinate ascent when optimizing AUC using the entire restricted reference (“whole AUC”) which represents the best achievable optimization, AUC using the subset of restricted reference (“partial AUC”) which represents AUC as used in practice, and the 3 other metrics mentioned above. The transcript assemblies constructed using the parameter vectors optimized using partial AUC as the objective recover more of the ground truth than those constructed using the default parameter vector. For the other metrics, we see that the transcript assemblies constructed with the parameter vectors found though optimization often showed a decrease in accuracy with respect to the transcripts constructed using the default parameter vector. These metrics are being lead astray by not incorporating the knowledge contained in the reference transcriptome.

**Figure 11:**
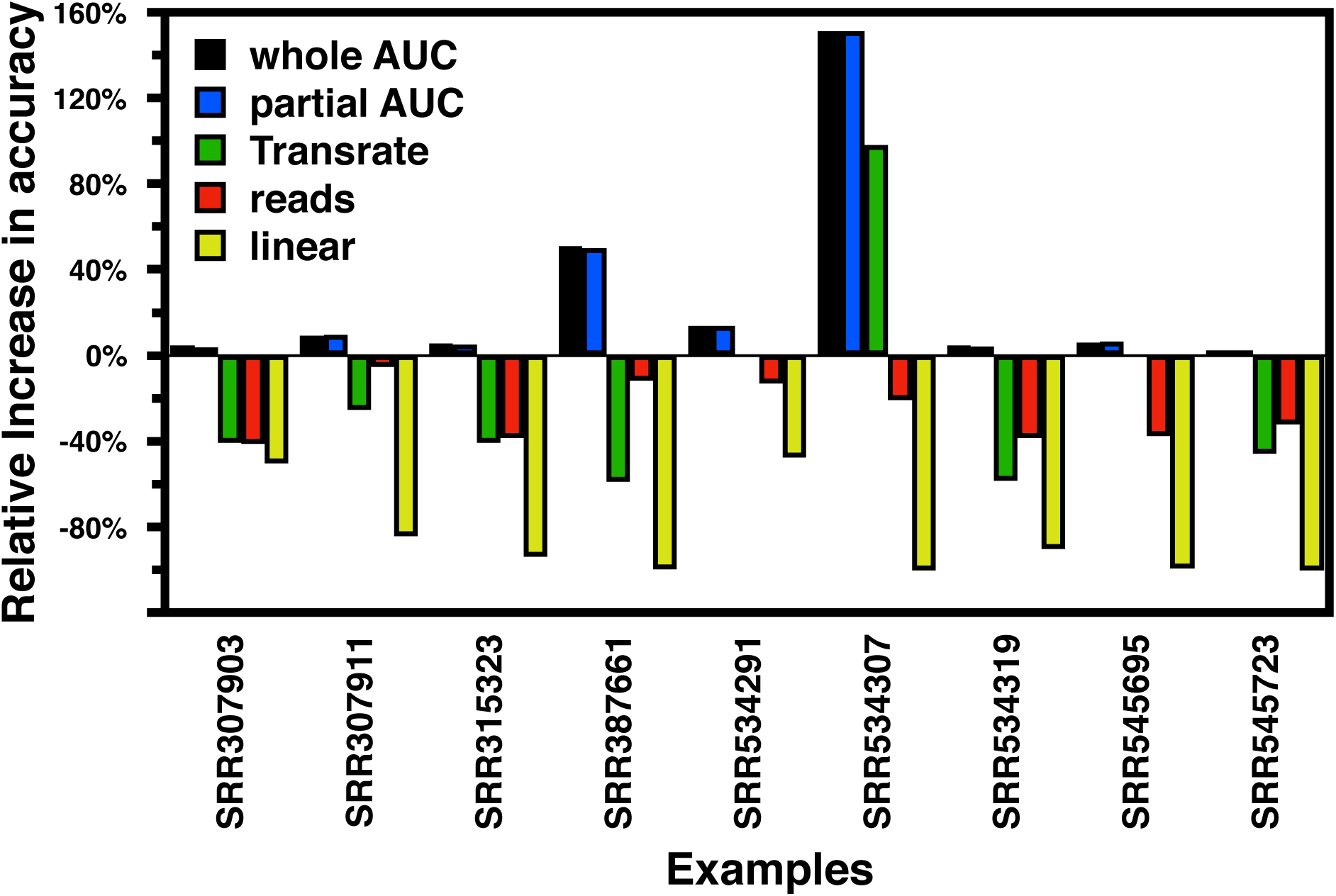
Relative increase in accuracy over default of using various transcript assembly metrics for coordinate ascent. The bars represent the difference in accuracy between the optimal parameter choice for each metric and the defaults, normalized by the default accuracy for each of the examples from `ENCODE10`.

An ideal test for how much predictive power is lost by optimizing AUC as opposed to other metrics would be to use a fully simulated RNA-seq sample where we know all of the transcripts that are present We choose to use experiments that more closely resemble biological samples in that they include extraneous sequencing reads that comes from amplification, sequencing, or assembly errors.

## 4. Conclusions

Our results show that sample-specific parameter vectors are important for developing any genomic pipeline that includes transcriptome assembly as a step. We begin to answer the question of how to produce transcriptome assemblies effectively for any input without sacrificing quality or expanding manpower. This is done using a combination of parameter tuning though exploration using coordinate ascent and the established method of parameter advising. Two key insights that made this merger viable and distinguish transcriptome assembly from other domains are: (1) because the parameter landscape likely has few local maxima, coordinate ascent rather than exhaustive enumeration can be used to find advisor sets, and (2) that a small, but representative, number of training examples is sufficient to provide a large increase in AUC over the default parameter setting.

The coordinate ascent procedure is a very useful means for finding parameter settings that increase the AUC of predicted transcriptomes. The best results can generally be achieved by simply following this procedure for each new input, although at a high running time. Our implementation of coordinate ascent does not allow steps to be taken in multiple directions at once. It is thus difficult to efficiently parallelize this process. In other words, coordinate ascent finds more accurate parameter vectors at the cost of large computational time. Instead, we have developed a method, which can be reapplied to any domain that has the same parameter behavior we observed, to find an advising set that is as diverse as the training examples used.

One drawback of using a single example to find each parameter vector in the set is that coordinate ascent is likely to overfit to the training examples. We have shown that even with this potential issue, we are able to improve the quality of the transcripts produced according to the AUC measure.

All of the results that we have shown assume that all transcripts in a sequencing sample can be assembled with a single choice of parameter vector. While this is an assumption made by most transcript assemblers, it may not be true in practice. One extension of this work is to provide *transcript-level* parameter choices to help improve the assembly quality. This would require the adaptation of AUC, or some other metric that could work well for parameter optimization, to provide transcript-level assessment.

The method we have described automates the task of parameter selection for `Scallop` and `StringTie`, and improves the quality of produced transcriptomes according to the area under the curve measure comparing the output to the reference transcriptome database. As we have shown, this measure works better at the task of parameter optimization than the existing *de novo* measures. But our results also show that there is room to improve this measure since in Figure 11 partial AUC is never able to recover all of the “novelty” present in the sample. The unknown transcripts can sometimes be the most interesting, and this measure of accuracy penalizes novelty by definition, and the assembler may not recover this unknown transcript if it is highly divergent from those we already know.

An alternate explanation of the results in Section 3.5 are that the assemblies produced by the parameter vectors found by optimizing the *de novo* metrics to be optimal are actually more realistic than those found for AUC. For this to be true this would mean there is a large number of false positives found when constructing a transcript assembly using the default parameter vector. Even though there is some debate over the completeness of the reference transcriptome [26, 27], this seems to be unlikely. In the future, it would be ideal to find some new measure that is some hybrid between AUC and *de novo* assessment that could both use current knowledge but still be able to detect novel transcripts. When this is available, it can replace AUC as the measure for constructing and using advising for transcript assembly.

## Acknowledgements

The authors would like to thank the members of the Kingsford Group for their helpful comments through out this project, in particular Mingfu Shao, Jonathan King, Heewook Lee, Guillaume Marçais, Prashant Pandey, and Minh Hoàng.

## Footnotes

1 https://github.com/Kingsford-Group/scallopadvising

2 https://github.com/gpertea/gffcompare

3 https://github.com/Kingsford-Group/rnaseqtools

## References

[1] Dan DeBlasio and John Kececioglu. Parameter advising for multiple sequence alignment, volume 26 of Computational Biology series. Springer International Publishing, Cham, Switzerland, 2017.

[2] C. Trapnell, B.A. Williams, G. Pertea, A. Mortazavi, G. Kwan, M.J. Van Baren, S.L. Salzberg, B.J. Wold, and L. Pachter. Transcript assembly and quantification by RNA-Seq reveals unannotated transcripts and isoform switching during cell differentiation. Natue Biotechnology, 28(5):511–515, 2010.

[3] Mihaela Pertea, Geo M Pertea, Corina M Antonescu, Tsung-Cheng Chang, Joshua T Mendell, and Steven L Salzberg. StringTie enables improved reconstruction of a transcriptome from RNA-Seq reads. Nature Biotechnology, 33:290–295, February 2015.

[4] Juntao Liu, Ting Yu, Tao Jiang, and Guojun Li. TransComb: genome-guided transcriptome assembly via combing junctions in splicing graphs. Genome Biology, 17(1):213, October 2016.

[5] Mingfu Shao and Carl Kingsford. Accurate assembly of transcripts through phase-preserving graph decomposition. Nature Biotechnology, 35:1167–1169, 2017.

[6] Daehwan Kim, Ben Langmead, and Steven L Salzberg. HISAT: a fast spliced aligner with low memory requirements. Nature Methods, 12:357–360, March 2015.

[7] Alexander Dobin, Carrie A. Davis, Felix Schlesinger, Jorg Drenkow, Chris Zaleski, Sonali Jha, Philippe Batut, Mark Chaisson, and Thomas R. Gingeras. STAR: ultrafast universal RNA-Seq aligner. Bioinformatics, 29(1):15–21, 2013.

[8] Daehwan Kim, Geo Pertea, Cole Trapnell, Harold Pimentel, Ryan Kelley, and Steven L. Salzberg. TopHat2: accurate alignment of transcriptomes in the presence of insertions, deletions and gene fusions. Genome Biology, 14(4):R36, April 2013.

[9] K.F. Au, H. Jiang, L. Lin, Y. Xing, and W.H. Wong. Detection of splice junctions from paired-end RNA-Seq data by splicemap. Nucleic Acids Research, 38(14):4570–4578, 2010.

[10] Rob Patro, Geet Duggal, Michael I Love, Rafael A Irizarry, and Carl Kingsford. Salmon provides fast and bias-aware quantification of transcript expression. Nature Methods, 14(4):417–419, 2017.

[11] Nicolas L Bray, Harold Pimentel, Páll Melsted, and Lior Pachter. Near-optimal probabilistic rna-seq quantification. Nature Biotechnology, 34, April 2016.

[12] Michael I Love, Wolfgang Huber, and Simon Anders. Moderated estimation of fold change and dispersion for RNA-Seq data with deseq2. Genome Biology, 15(12):550, 2014.

[13] Alyssa C Frazee, Geo Pertea, Andrew E Jaffe, Ben Langmead, Steven L Salzberg, and Jeffrey T Leek. Flexible isoform-level differential expression analysis with ballgown. Nature Biotechnology, 33:243—-246, March 2015.

[14] Lin Xu, Frank Hutter, Holger H Hoos, and Kevin Leyton-Brown. SATzilla: portfolio-based algorithm selection for SAT. J. Artif. Intel. Res., 32:565–606, 2008.

[15] Frank Hutter, Holger H. Hoos, Kevin Leyton-Brown, and Thomas Stützle. ParamILS: an automatic algorithm configuration framework. J. Artif. Intel. Res., 36(1):267–306, September 2009.

[16] Rayan Chikhi and Paul Medvedev. Informed and automated k-mer size selection for genome assembly. Bioinformatics, 30(1):31–37, June 2013.

[17] Randal S. Olson, Nathan Bartley, Ryan J. Urbanowicz, and Jason H. Moore. Evaluation of a tree-based pipeline optimization tool for automating data science. In Proceedings of the Genetic and Evolutionary Computation Conference 2016, pages 485–492, 2016.

[18] Jasper Snoek, Hugo Larochelle, and Ryan P Adams. Practical bayesian optimization of machine learning algorithms. In Proceedings of Advances in Neural Information Processing Systems, pages 2951–5959, 2012.

[19] Augustin Cauchy. Méthode générale pour la résolution des systémes d’équations simultanées. Comptes Rendus Hebdomadaire des Séances de l’Academie des Sciences, 25(536):536–538, 1847.

[20] S. Kirkpatrick, C. D. Gelatt, and M. P. Vecchi. Optimization by simulated annealing. Science, 220(4598):671–680, May 1983.

[21] Willard I. Zangwill. Nonlinear Programming: A Unified Approach. Prentice-Hall International, Englewood Cliffs, N.J, 1969.

[22] The ENCODE Project Consortium. An integrated encyclopedia of DNA elements in the human genome. Nature, 489:57–74, August 2012.

[23] Rasko Leinonen, Hideaki Sugawara, and Shumway, M. on behalf of the International Nucleotide Sequence Database Collaboration. The sequence read archive. Nucleic Acids Research, 39:D19–D21, November 2010.

[24] Dan DeBlasio and John Kececioglu. Learning parameter-advising sets for multiple sequence alignment. IEEE/ACM Transactions on Computational Biology and Bioinformatics, 14(5):1028–1041, 2017.

[25] Richard Smith-Unna, Chris Boursnell, Rob Patro, Julian M Hibberd, and Steven Kelly. TransRate: reference-free quality assessment of de novo transcriptome assemblies. Genome Research, 26(8):1134–1144, August 2016.

[26] Mihaela Pertea, Alaina Shumate, Geo Pertea, Ales Varabyou, Florian P Breitwieser, Yu-Chi Chang, Anil K Madugundu, Akhilesh Pandey, and Steven L Salzberg. CHESS: a new human gene catalog curated from thousands of large-scale RNA sequencing experiments reveals extensive transcriptional noise. Genome Biology, 19(1):208, November 2018.

[27] Irwin Jungreis, Michael L. Tress, Jonathan Mudge, Cristina Sisu, Toby Hunt, Rory Johnson, Barbara Uszczynska-Ratajczak, Julien Lagarde, James Wright, Paul Muir, Mark Gerstein, Roderic Guigo, Manolis Kellis, Adam Frankish, Paul Flicek, and The GENCODE Consortium. Nearly all new protein-coding predictions in the CHESS database are not protein-coding. bioRxiv, 360602, 2018.

