## Supplemental Tables for "More accurate transcript assembly via parameter advising"

### Towards building an automated bioinformatician: more accurate, large-scale genomic discovery via parameter advising

This document contains additional experimental information, information about the versions and command line arguments for the software used, as well as the experiment identifiers for the samples from `ENCODE` and `SRA` used in the manuscript.

Table 1: Optimal parameter vectors found by coordinate ascent

| ID | Experiment | Aligner | ICC | NE | BG | EL | FL | MQ | NH | SBH | SG | SL | SO | TLB | TLI | UM | US | 1 | 2 | 4 | 8 |
|---|---|---|---|---|---|---|---|---|---|---|---|---|---|---|---|---|---|---|---|---|---|
| 1 | SRR545723 | `TopHat` | 2.00 | 1000 | 1359 | 20 | 3 | 11 | 20 | 1 | 39 | 11 | 1.50 | 454 | 8 | false | false | X | | X | X |
| 2 | SRR307911 | `TopHat` | 2.00 | 1000 | 1032 | 20 | 2 | 11 | 21 | 1 | 52 | 15 | 1.50 | 432 | 19 | false | false | | | | |
| 3 | SRR534291 | `TopHat` | 0.75 | 51 | 785 | 25 | 3 | 1 | 9 | 1 | 121 | 16 | 1.51 | 503 | 8 | false | false | | | | X |
| 4 | SRR387661 | `TopHat` | 0.22 | 1000 | 999 | 20 | 3 | 11 | 20 | 1 | 53 | 15 | 1.50 | 429 | 19 | false | false | | X | | |
| 5 | SRR315334 | `TopHat` | 1.50 | 1000 | 1425 | 20 | 3 | 11 | 20 | 1 | 50 | 16 | 1.50 | 528 | 1 | false | false | | | | |
| 6 | SRR307903 | `TopHat` | 1.61 | 250 | 733 | 2 | 0 | 11 | 2 | 1 | 23 | 26 | 0.75 | 521 | 2 | false | false | | | | |
| 7 | SRR545695 | `TopHat` | 2.00 | 1000 | 852 | 20 | 2 | 2 | 104 | 1 | 5 | 7 | 1.50 | 432 | 19 | false | false | | | | |
| 8 | SRR534319 | `TopHat` | 2.27 | 1000 | 306 | 20 | 0 | 1 | 20 | 1 | 7 | 11 | 1.50 | 346 | 21 | false | false | | | | |
| 9 | SRR315323 | `TopHat` | 0.22 | 205 | 945 | 20 | 0 | 2 | 20 | 1 | 39 | 15 | 1.16 | 409 | 8 | true | true | | | | |
| 10 | SRR534307 | `TopHat` | 2.22 | 1000 | 78 | 32 | 3 | 11 | 16 | 2 | 0 | 32 | 1.50 | 512 | 8 | false | false | | X | | |
| 11 | SRR545723 | `HISAT` | 1.75 | 11000 | 628 | 20 | 3 | 11 | 20 | 2 | 8 | 7 | 1.50 | 408 | 8 | false | false | | | | |
| 12 | SRR307911 | `HISAT` | 99.66 | 1000 | 1828 | 3 | 5 | 11 | 3 | 1 | 43 | 15 | 0.17 | 392 | 8 | false | false | | | | |
| 13 | SRR534291 | `HISAT` | 1.00 | 11000 | 851 | 22 | 5 | 11 | 3 | 1 | 120 | 3 | 0.17 | 267 | 30 | false | false | | | | |
| 14 | SRR387661 | `HISAT` | 1.05 | 454 | 130 | 26 | 4 | 11 | 38 | 1 | 23 | 0 | 0.17 | 880 | 0 | false | false | | | | X |
| 15 | SRR315334 | `HISAT` | 0.00 | 282 | 870 | 20 | 4 | 11 | 1 | 1 | 48 | 8 | 1.28 | 307 | 21 | false | false | | | | |
| 16 | SRR307903 | `HISAT` | 1.42 | 168 | 745 | 9 | 5 | 11 | 2 | 1 | 23 | 6 | 0.17 | 396 | 0 | false | false | | | | |
| 17 | SRR545695 | `HISAT` | 5.56 | 1000 | 778 | 26 | 2 | 11 | 25 | 2 | 0 | 0 | 1.50 | 360 | 8 | false | false | | | | X |
| 18 | SRR534319 | `HISAT` | 2.00 | 1000 | 1669 | 20 | 0 | 11 | 81 | 1 | 4 | 7 | 1.50 | 150 | 73 | false | false | | | | |
| 19 | SRR315323 | `HISAT` | 2.00 | 685 | 782 | 15 | 0 | 11 | 14 | 1 | 40 | 7 | 1.50 | 180 | 50 | false | false | | | | |
| 20 | SRR534307 | `HISAT` | 2.00 | 146 | 309 | 41 | 8 | 11 | 10 | 3 | 2 | 0 | 1.17 | 370 | 4 | false | false | | | X | X |
| 21 | SRR545723 | `STAR` | 2.00 | 11000 | 455 | 22 | 6 | 11 | 3 | 1 | 3 | 21 | 1.26 | 252 | 16 | false | false | | | | |
| 22 | SRR307911 | `STAR` | 0.22 | 1000 | 490 | 20 | 3 | 11 | 8 | 1 | 53 | 26 | 0.50 | 289 | 24 | false | false | | | X | |
| 23 | SRR534291 | `STAR` | 0.22 | 11000 | 481 | 24 | 0 | 2 | 11 | 1 | 123 | 26 | 0.24 | 352 | 8 | false | false | | | | |
| 24 | SRR387661 | `STAR` | 2.00 | 1000 | 734 | 20 | 4 | 11 | 35 | 2 | 42 | 29 | 1.50 | 411 | 1 | false | false | | | | |
| 25 | SRR315334 | `STAR` | 0.50 | 1000 | 1070 | 8 | 0 | 11 | 8 | 1 | 41 | 29 | 0.75 | 267 | 33 | false | false | | | | X |
| 26 | SRR307903 | `STAR` | 2.00 | 1000 | 976 | 8 | 5 | 11 | 20 | 1 | 35 | 35 | 0.17 | 433 | 8 | false | false | | | | |
| 27 | SRR545695 | `STAR` | 2.00 | 1000 | 388 | 20 | 8 | 11 | 89 | 2 | 20 | 23 | 1.50 | 307 | 19 | false | false | | | | |
| 28 | SRR534319 | `STAR` | 2.00 | 800 | 1574 | 17 | 4 | 2 | 20 | 1 | 8 | 18 | 1.50 | 307 | 4 | false | false | | | | X |
| 29 | SRR315323 | `STAR` | 2.00 | 11000 | 1043 | 20 | 3 | 7 | 20 | 1 | 33 | 35 | 1.50 | 150 | 50 | true | true | | | | |
| 30 | SRR534307 | `STAR` | 1.58 | 292 | 54 | 20 | 0 | 11 | 9 | 3 | 0 | 29 | 1.50 | 320 | 8 | false | false | | | X | X |
| | Default | | 2.00 | 1,000 | 50 | 20 | 3 | 1 | 20 | 1 | 3 | 15 | 1.50 | 150 | 50 | false | false | | | | |
| | Initial step size | | 10.00 | 10,000 | 100 | 100 | 10 | 10 | 100 | 10 | 10 | 20 | 10.00 | 500 | 100 | n/a | n/a | | | | |

Table 2: Increase in AUC for examples in `ENCODE10`. Percentages in alternating rows show the increase over the default.

| | `HISAT` | | | `STAR` | | | `TopHat` | | |
|---|---|---|---|---|---|---|---|---|---|
| Experiment | Default | CA | LOO | Default | CA | LOO | Default | CA | LOO |
| SRR307903 | 589.271 | 612.017 | 604.268 | 528.307 | 556.867 | 548.746 | 448.498 | 496.242 | 483.381 |
| | | 3.86% | 2.54% | | 5.4% | 3.86% | | 10.64% | 7.77% |
| SRR307911 | 503.460 | 549.553 | 544.157 | 477.174 | 519.346 | 514.035 | 387.947 | 446.299 | 438.225 |
| | | 9.15% | 8.08% | | 8.83% | 7.72% | | 15.04% | 12.96% |
| SRR315323 | 389.409 | 409.204 | 408.548 | 340.764 | 351.396 | 349.285 | 283.773 | 308.121 | 305.539 |
| | | 5.08% | 4.91% | | 3.12% | 2.5% | | 8.58% | 7.67% |
| SRR315334 | 549.081 | 579.641 | 573.181 | 501.790 | 532.407 | 525.885 | 413.393 | 472.124 | 466.392 |
| | | 5.56% | 4.38% | | 6.1% | 4.8% | | 14.2% | 12.82% |
| SRR387661 | 199.230 | 299.722 | 277.117 | 464.403 | 493.931 | 487.166 | 168.373 | 458.468 | 453.251 |
| | | 50.44% | 39.09% | | 6.35% | 4.9% | | 172.29% | 169.19% |
| SRR534291 | 469.952 | 533.842 | 522.227 | 432.317 | 496.602 | 481.104 | 388.038 | 460.242 | 440.755 |
| | | 13.59% | 11.12% | | 14.86% | 11.28% | | 18.6% | 13.58% |
| SRR534307 | 293.485 | 734.992 | 710.496 | 639.449 | 716.494 | 659.162 | 638.106 | 692.815 | 678.095 |
| | | 150.43% | 142.08% | | 12.04% | 3.08% | | 8.57% | 6.26% |
| SRR534319 | 303.001 | 313.419 | 308.410 | 257.493 | 267.573 | 260.004 | 244.602 | 267.438 | 257.182 |
| | | 3.43% | 1.78% | | 3.91% | 0.97% | | 9.33% | 5.14% |
| SRR545695 | 370.929 | 393.646 | 385.311 | 338.668 | 349.860 | 344.742 | 152.441 | 346.034 | 337.442 |
| | | 6.12% | 3.87% | | 3.3% | 1.79% | | 126.99% | 121.35% |
| SRR545723 | 537.776 | 551.086 | 548.686 | 525.978 | 533.369 | 524.506 | 458.757 | 492.322 | 485.531 |
| | | 2.47% | 2.02% | | 1.4% | -0.27% | | 7.31% | 5.83% |

Table 3: Command line options and software versions used

| Software | Version | Command |
|---|---|---|
| Scallop | v0.10.2 | scallop <options> -min_transcript_coverage 0\ |
| | | -i <alignment> -o <assembly> |
| alternate* | | scallop_run_with_config.pl <output_dir> <config> <alignment> |
| StringTie | v1.3.2d | stringtie <options> -c 0.001 -i <alignment> -o <assembly> |
| alternate* | | stringtie_run_with_config.pl <output_dir> <config> <alignment> |
| GFFCompare | v0.10.1 | gffcompare -r <reference> -o <comparison_prefix> <assembly> |
| GTFCuff | v1.0.1 | gtfcuff auc <comparison_tmap> <number_of_transcripts> |
| Tophat2† | v2.1.1 | tophat2 -p 6 <index> <fastq1> <fastq2> |
| STAR† | v2.5.2a | STAR --outSAMstrandField <intron_motif> --chimSegmentMin 20 \ |
| | | --runThreadN 6 --genomeDir <index> --readFilesIn <fastq1> <fastq2> |
| HISAT† | v2.0.4 | hisat2 -p 6 -x <index> -1 <fastq1> -2 <fastq2> |

* this command runs Scallop or StringTie from the configuration as well as GFFCompare and GTFCuff. These scripts are available in the Github repository (https://github.com/Kingsford-Group/scallopadvising).

† alignments used were acquired from the authors of Scallop (Shao and Kingsford 2017) the parameters here are those reported for their tests.

Table 4: Optimal parameter vectors for `StringTie` found by coordinate ascent

| ID | Experiment | Aligner | M | a | f | g | j | m | t | u |
|---|---|---|---|---|---|---|---|---|---|---|
| 1 | SRR307903 | `HISAT` | 0.13 | 14 | 0 | 52 | 1 | 468 | false | false |
| 2 | SRR307911 | `HISAT` | 0.11 | 13 | 0 | 54 | 1 | 444 | false | false |
| 3 | SRR315323 | `HISAT` | 0.10 | 11 | 0 | 37 | 1 | 389 | false | false |
| 4 | SRR315334 | `HISAT` | 3.03 | 10 | 0 | 58 | 2 | 446 | false | false |
| 5 | SRR387661 | `HISAT` | 0.93 | 10 | 0 | 0 | 3 | 543 | false | false |
| 6 | SRR534291 | `HISAT` | 50.92 | 20 | 0.01 | 133 | 0 | 483 | false | false |
| 7 | SRR534307 | `HISAT` | 3.74 | 13 | 0 | 4 | 6 | 474 | false | false |
| 8 | SRR534319 | `HISAT` | 0.31 | 6 | 0.23 | 15 | 3 | 375 | false | true |
| 9 | SRR545695 | `HISAT` | 0.06 | 10 | 0.15 | 11 | 3 | 352 | false | false |
| 10 | SRR545723 | `HISAT` | 0.11 | 0 | 0 | 11 | 2 | 634 | false | false |
| 11 | SRR307903 | `STAR` | 0.34 | 6 | 0 | 21 | 2 | 464 | false | false |
| 12 | SRR307911 | `STAR` | 0.23 | 11 | 0.08 | 71 | 1 | 416 | false | false |
| 13 | SRR315323 | `STAR` | 50.92 | 11 | 0.19 | 54 | 0 | 382 | false | false |
| 14 | SRR315334 | `STAR` | 0.57 | 10 | 0 | 43 | 2 | 459 | false | false |
| 15 | SRR387661 | `STAR` | 50.92 | 13 | 0.1 | 77 | 4 | 408 | false | false |
| 16 | SRR534291 | `STAR` | 1.01 | 12 | 0 | 58 | 1 | 293 | false | false |
| 17 | SRR534307 | `STAR` | 50.89 | 8 | 0 | 1 | 7 | 596 | false | false |
| 18 | SRR534319 | `STAR` | 0.28 | 9 | 0.27 | 39 | 3 | 406 | false | false |
| 19 | SRR545695 | `STAR` | 0.31 | 10 | 0.19 | 62 | 4 | 456 | false | false |
| 20 | SRR545723 | `STAR` | 0.46 | 10 | 0.01 | 3 | 1 | 551 | false | false |
| 21 | SRR307903 | `TopHat` | 0.06 | 10 | 0 | 64 | 1 | 575 | false | false |
| 22 | SRR307911 | `TopHat` | 0.08 | 8 | 0 | 53 | 1 | 561 | false | false |
| 23 | SRR315323 | `TopHat` | 0.29 | 1 | 0 | 40 | 0 | 536 | false | false |
| 24 | SRR315334 | `TopHat` | 0.07 | 7 | 0 | 38 | 2 | 530 | false | false |
| 25 | SRR387661 | `TopHat` | 0.21 | 6 | 0 | 9 | 3 | 579 | false | false |
| 26 | SRR534291 | `TopHat` | 12.79 | 10 | 0 | 112 | 1 | 492 | false | false |
| 27 | SRR534307 | `TopHat` | 0.12 | 7 | 0 | 2 | 5 | 593 | false | false |
| 28 | SRR534319 | `TopHat` | 0.18 | 5 | 0.25 | 39 | 2 | 370 | false | false |
| 29 | SRR545695 | `TopHat` | 0.23 | 8 | 0.13 | 14 | 3 | 407 | false | false |
| 30 | SRR545723 | `TopHat` | 0.18 | 3 | 0 | 61 | 1 | 778 | false | false |
| | Default | | 0.95 | 10 | 0.1 | 50 | 1 | 10 | false | false |
| | Initial Step Size | | 50 | 1,000 | 50 | 5,000 | 100 | 10,000 | n/a | n/a |

M — fraction of bundle allowed to be covered by multi-hit reads
a — minimum anchor length for junctions
f — minimum isoform fraction
g — gap between read mappings triggering a new bundle
j — minimum junction coverage
m — minimum assembled transcript length
t — disable trimming of predicted transcripts based on coverage
u — no multi-mapping correction

Table 5: Increase in `StringTie` AUC for examples in `ENCODE10`

| | | `HISAT` | | | `STAR` | | | `TopHat` | |
|---|---|---|---|---|---|---|---|---|---|
| Experiment | Default | CA | LOO | Default | CA | LOO | Default | CA | LOO |
| SRR307903 | 465.718 | 512.245 | 511.875 | 392.436 | 413.918 | 409.366 | 362.760 | 419.973 | 415.095 |
| | | 9.99% | 9.91% | | 5.47% | 4.31% | | 15.77% | 14.42% |
| SRR307911 | 410.998 | 434.589 | 425.456 | 358.409 | 369.190 | 366.879 | 310.537 | 365.481 | 358.772 |
| | | 5.73% | 3.51% | | 3% | 2.36% | | 17.69% | 15.53% |
| SRR315323 | 313.313 | 326.525 | 319.800 | 267.734 | 272.503 | 259.881 | 234.834 | 257.797 | 253.212 |
| | | 4.21% | 2.07% | | 1.78% | -2.93% | | 9.77% | 7.82% |
| SRR315334 | 420.644 | 470.429 | 463.242 | 354.466 | 383.854 | 375.994 | 328.576 | 395.386 | 385.928 |
| | | 11.83% | 10.12% | | 8.29% | 6.07% | | 20.33% | 17.45% |
| SRR387661 | 367.649 | 411.868 | 404.578 | 318.456 | 342.697 | 339.809 | 312.360 | 363.039 | 356.312 |
| | | 12.02% | 10.04% | | 7.61% | 6.7% | | 16.22% | 14.07% |
| SRR534291 | 400.711 | 431.269 | 428.479 | 344.799 | 361.791 | 356.259 | 338.306 | 379.263 | 381.298 |
| | | 7.62% | 6.92% | | 4.92% | 3.32% | | 12.1% | 12.7% |
| SRR534307 | 489.710 | 585.195 | 560.411 | 449.131 | 517.549 | 506.824 | 472.057 | 552.523 | 545.428 |
| | | 19.49% | 14.43% | | 15.23% | 12.84% | | 17.04% | 15.54% |
| SRR534319 | 197.555 | 224.598 | 217.561 | 165.125 | 186.765 | 182.049 | 171.739 | 193.674 | 188.326 |
| | | 13.68% | 10.12% | | 13.1% | 10.24% | | 12.77% | 9.65% |
| SRR545695 | 264.356 | 297.862 | 292.469 | 230.141 | 265.741 | 258.789 | 235.576 | 264.462 | 256.935 |
| | | 12.67% | 10.63% | | 15.46% | 12.44% | | 12.26% | 9.06% |
| SRR545723 | 430.262 | 467.512 | 458.859 | 404.491 | 429.514 | 416.254 | 370.397 | 416.176 | 406.089 |
| | | 8.65% | 6.64% | | 6.18% | 2.9% | | 12.35% | 9.63% |

CA — Parameter vector found using coordinate ascent
LOO — Advising using leave-one-out set, the 18 parameters with different experiment and aligner

Table 6: `ENCODE10` Samples

| | | | | |
|---|---|---|---|---|
| SRR307903 | SRR307911 | SRR315323 | SRR315334 | SRR387661 |
| SRR534291 | SRR534307 | SRR534319 | SRR545695 | SRR545723 |

Table 7: `ENCODE65` Samples

| | | | | |
|---|---|---|---|---|
| ENCFF000CXQ | ENCFF000DBO | ENCFF000DDA | ENCFF000DGH | ENCFF000DLJ |
| ENCFF000DNB | ENCFF000DPP | ENCFF000DRN | ENCFF000DTX | ENCFF000DYK |
| ENCFF000EAF | ENCFF000EBB | ENCFF000ECJ | ENCFF000EEW | ENCFF000EFQ |
| ENCFF001RRX | ENCFF001RSM | ENCFF001RWH | ENCFF001RWL | ENCFF002DKQ |
| ENCFF002DKV | ENCFF019PWN | ENCFF020BHL | ENCFF020JKV | ENCFF023MNP |
| ENCFF024INL | ENCFF031DGP | ENCFF040LQY | ENCFF044SJL | ENCFF046HDL |
| ENCFF050RHK | ENCFF052GIA | ENCFF054RXE | ENCFF068GSV | ENCFF075XPF |
| ENCFF084JYA | ENCFF088RGO | ENCFF095MSA | ENCFF098AZJ | ENCFF100BIM |
| ENCFF101NVY | ENCFF126PQC | ENCFF130QNU | ENCFF134DLO | ENCFF135DSP |
| ENCFF136KRX | ENCFF142WMA | ENCFF166HWB | ENCFF172AWE | ENCFF176HWN |
| ENCFF216PVC | ENCFF227NHT | ENCFF233ICM | ENCFF240IBI | ENCFF255JTL |
| ENCFF257GEY | ENCFF296GRX | ENCFF306YQS | ENCFF316NIV | ENCFF386PAW |
| ENCFF448GEE | ENCFF455XNT | ENCFF537YYU | ENCFF685IGY | ENCFF710AVC |

Table 8: SRA Samples

| | | | | | |
|---|---|---|---|---|---|
| DRR021368 | DRR021561 | DRR021562 | DRR023030 | ERR030872 | ERR030873 |
| ERR030874 | ERR030875 | ERR030876 | ERR030877 | ERR030878 | ERR030879 |
| ERR030886 | ERR030887 | ERR188025 | ERR188029 | ERR188050 | ERR188052 |
| ERR188054 | ERR188055 | ERR188056 | ERR188057 | ERR188070 | ERR188072 |
| ERR188073 | ERR188074 | ERR188077 | ERR188084 | ERR188089 | ERR188091 |
| ERR188093 | ERR188097 | ERR188111 | ERR188112 | ERR188114 | ERR188122 |
| ERR188128 | ERR188132 | ERR188135 | ERR188137 | ERR188140 | ERR188141 |
| ERR188144 | ERR188145 | ERR188152 | ERR188153 | ERR188154 | ERR188156 |
| ERR188157 | ERR188158 | ERR188164 | ERR188169 | ERR188194 | ERR188198 |
| ERR188237 | ERR188261 | ERR188264 | ERR188265 | ERR188269 | ERR188270 |
| ERR188272 | ERR188295 | ERR188318 | ERR188330 | ERR188337 | ERR188353 |
| ERR188371 | ERR188372 | ERR188400 | ERR188403 | ERR188405 | ERR188408 |
| ERR188415 | ERR188418 | ERR188419 | ERR188430 | ERR188432 | ERR188459 |
| ERR188478 | ERR197921 | ERR197923 | ERR204868 | ERR204894 | ERR204897 |
| ERR204950 | ERR204993 | ERR208906 | ERR296101 | ERR296104 | ERR304481 |
| ERR304482 | ERR304483 | ERR304484 | ERR304486 | ERR304488 | ERR313180 |
| ERR313182 | ERR315338 | ERR315391 | ERR315448 | ERR315456 | ERR412874 |
| ERR412876 | ERR412877 | ERR532433 | ERR532434 | ERR532435 | ERR532436 |
| ERR532437 | ERR532438 | ERR532439 | ERR532440 | ERR532441 | ERR532442 |
| ERR532443 | ERR532444 | ERR532446 | ERR532447 | ERR532548 | ERR532550 |
| ERR532551 | ERR532560 | ERR532561 | ERR532562 | ERR532563 | ERR532565 |
| ERR532567 | ERR532568 | ERR532569 | ERR532593 | ERR532600 | ERR532601 |
| ERR532603 | ERR532604 | ERR579122 | ERR579127 | ERR579131 | ERR579136 |
| ERR579140 | ERR579146 | ERR579147 | ERR579148 | ERR658219 | ERR658221 |
| ERR661162 | ERR695500 | SRR1004852 | SRR1004853 | SRR1004854 | SRR1004855 |
| SRR1004856 | SRR1004857 | SRR1004858 | SRR1004859 | SRR1004860 | SRR1023010 |
| SRR1023011 | SRR1023790 | SRR1023792 | SRR1023793 | SRR1023794 | SRR1023795 |
| SRR1023796 | SRR1023797 | SRR1023798 | SRR1023799 | SRR1024048 | SRR1026880 |
| SRR1026881 | SRR1026882 | SRR1026883 | SRR1026884 | SRR1026886 | SRR1026887 |
| SRR1026888 | SRR1026889 | SRR1026908 | SRR1026920 | SRR1026921 | SRR1026922 |
| SRR1026924 | SRR1026925 | SRR1026926 | SRR1026927 | SRR1026928 | SRR1026929 |
| SRR1026939 | SRR1026940 | SRR1026941 | SRR1026942 | SRR1026943 | SRR1026944 |
| SRR1026945 | SRR1026948 | SRR1026949 | SRR1026956 | SRR1026959 | SRR1026989 |
| SRR1026990 | SRR1026991 | SRR1026993 | SRR1026996 | SRR1026997 | SRR1027000 |
| SRR1027002 | SRR1027004 | SRR1027006 | SRR1027007 | SRR1027010 | SRR1027181 |
| SRR1027182 | SRR1027183 | SRR1027184 | SRR1027185 | SRR1027186 | SRR1027188 |
| SRR1027732 | SRR1028343 | SRR1028344 | SRR1028345 | SRR1028346 | SRR1028347 |
| SRR1028348 | SRR1028349 | SRR1029601 | SRR1029602 | SRR1030493 | SRR1030495 |
| SRR1030496 | SRR1030497 | SRR1030499 | SRR1036041 | SRR1036050 | SRR1036054 |
| SRR1036085 | SRR1036086 | SRR1036087 | SRR1036089 | SRR1036105 | SRR1036106 |

Table 9: SRA Samples (continued)

| | | | | | |
|---|---|---|---|---|---|
| SRR1036107 | SRR1036109 | SRR1036114 | SRR1044667 | SRR1044669 | SRR1044670 |
| SRR1044672 | SRR1044673 | SRR1044675 | SRR1046364 | SRR1046683 | SRR1046686 |
| SRR1046797 | SRR1046816 | SRR1046823 | SRR1046825 | SRR1047863 | SRR1047864 |
| SRR1047865 | SRR1047867 | SRR1047868 | SRR1047869 | SRR1047872 | SRR1047873 |
| SRR1047874 | SRR1049826 | SRR1049827 | SRR1049828 | SRR1049829 | SRR1054287 |
| SRR1054290 | SRR1054291 | SRR1056401 | SRR1056402 | SRR1060754 | SRR1060757 |
| SRR1060766 | SRR1061339 | SRR1067911 | SRR1067912 | SRR1067913 | SRR1067914 |
| SRR1067915 | SRR1067916 | SRR1067917 | SRR1067918 | SRR1067919 | SRR1104902 |
| SRR1104903 | SRR1104905 | SRR1118401 | SRR1118402 | SRR1122357 | SRR1122358 |
| SRR1124845 | SRR1145838 | SRR1145839 | SRR1145840 | SRR1145842 | SRR1145843 |
| SRR1153278 | SRR1153279 | SRR1153284 | SRR1153470 | SRR1163130 | SRR1167725 |
| SRR1167726 | SRR1175193 | SRR1175194 | SRR1175195 | SRR1175196 | SRR1175198 |
| SRR1175199 | SRR1177667 | SRR1177668 | SRR1177669 | SRR1177692 | SRR1177694 |
| SRR1177695 | SRR1177698 | SRR1177699 | SRR1177705 | SRR1177708 | SRR1177709 |
| SRR1177720 | SRR1177721 | SRR1177722 | SRR1177723 | SRR1177724 | SRR1177725 |
| SRR1177726 | SRR1177727 | SRR1177728 | SRR1177729 | SRR1177740 | SRR1177741 |
| SRR1177742 | SRR1177743 | SRR1177744 | SRR1177745 | SRR1177746 | SRR1177747 |
| SRR1177749 | SRR1182250 | SRR1182251 | SRR1182256 | SRR1182257 | SRR1182262 |
| SRR1182263 | SRR1182264 | SRR1182265 | SRR1182266 | SRR1182267 | SRR1182268 |
| SRR1182269 | SRR1186053 | SRR1186601 | SRR1186603 | SRR1186606 | SRR1186610 |
| SRR1186611 | SRR1186612 | SRR1186613 | SRR1191667 | SRR1191668 | SRR1191876 |
| SRR1191914 | SRR1191953 | SRR1191954 | SRR1191990 | SRR1191991 | SRR1192321 |
| SRR1193014 | SRR1193015 | SRR1193016 | SRR1193100 | SRR1193101 | SRR1193102 |
| SRR1193103 | SRR1193104 | SRR1193105 | SRR1193106 | SRR1193198 | SRR1193638 |
| SRR1193639 | SRR1200501 | SRR1200503 | SRR1200507 | SRR1200511 | SRR1200515 |
| SRR1201619 | SRR1201621 | SRR1201623 | SRR1201625 | SRR1201629 | SRR1201631 |
| SRR1201641 | SRR1201662 | SRR1201664 | SRR1201666 | SRR1201670 | SRR1201672 |
| SRR1201673 | SRR1201675 | SRR1201676 | SRR1201677 | SRR1201706 | SRR1201707 |
| SRR1201708 | SRR1201721 | SRR1201723 | SRR1201725 | SRR1201727 | SRR1201743 |
| SRR1201745 | SRR1201750 | SRR1201754 | SRR1201761 | SRR1201763 | SRR1201765 |
| SRR1201767 | SRR1201768 | SRR1201792 | SRR1201794 | SRR1201798 | SRR1201814 |
| SRR1201815 | SRR1201816 | SRR1201817 | SRR1201818 | SRR1201819 | SRR1201820 |
| SRR1201828 | SRR1201829 | SRR1201830 | SRR1201831 | SRR1201832 | SRR1201833 |
| SRR1201982 | SRR1201997 | SRR1201998 | SRR1202093 | SRR1202289 | SRR1202564 |
| SRR1202651 | SRR1202846 | SRR1202847 | SRR1203132 | SRR1206036 | SRR1206038 |
| SRR1206039 | SRR1206040 | SRR1206042 | SRR1206043 | SRR1206044 | SRR1206045 |
| SRR1220685 | SRR1220686 | SRR1220687 | SRR1220688 | SRR1220689 | SRR1220690 |
| SRR1220691 | SRR1220692 | SRR1220693 | SRR1220694 | SRR1220695 | SRR1220696 |
| SRR1220697 | SRR1220698 | SRR1232214 | SRR1232247 | SRR1232253 | SRR1237975 |
| SRR1237976 | SRR1237978 | SRR1237979 | SRR1237990 | SRR1237991 | SRR1237992 |

Table 10: SRA Samples (continued)

| | | | | | |
|---|---|---|---|---|---|
| SRR1237993 | SRR1237994 | SRR1238549 | SRR1239514 | SRR1239515 | SRR1239516 |
| SRR1239517 | SRR1240808 | SRR1240809 | SRR1248259 | SRR1248260 | SRR1248261 |
| SRR1258220 | SRR1261168 | SRR1261169 | SRR1261170 | SRR1264914 | SRR1265054 |
| SRR1266980 | SRR1266981 | SRR1268166 | SRR1268169 | SRR1268173 | SRR1269755 |
| SRR1270780 | SRR1270799 | SRR1270835 | SRR1270836 | SRR1273699 | SRR1273700 |
| SRR1273701 | SRR1273702 | SRR1273703 | SRR1273704 | SRR1273706 | SRR1273707 |
| SRR1273708 | SRR1273709 | SRR1275413 | SRR1283008 | SRR1286920 | SRR1286923 |
| SRR1286924 | SRR1286925 | SRR1286927 | SRR1286928 | SRR1286929 | SRR1287012 |
| SRR1292695 | SRR1292696 | SRR1292697 | SRR1296120 | SRR1296121 | SRR1296125 |
| SRR1296127 | SRR1296128 | SRR1296129 | SRR1296136 | SRR1296138 | SRR1296142 |
| SRR1297311 | SRR1297312 | SRR1297317 | SRR1299472 | SRR1299473 | SRR1299474 |
| SRR1299475 | SRR1313067 | SRR1313068 | SRR1313069 | SRR1313070 | SRR1313071 |
| SRR1313072 | SRR1313073 | SRR1313074 | SRR1313075 | SRR1313076 | SRR1313079 |
| SRR1313080 | SRR1313086 | SRR1313088 | SRR1313090 | SRR1313091 | SRR1313092 |
| SRR1313093 | SRR1313094 | SRR1313095 | SRR1313096 | SRR1313097 | SRR1313098 |
| SRR1313102 | SRR1313114 | SRR1313115 | SRR1313119 | SRR1313130 | SRR1313131 |
| SRR1313133 | SRR1313134 | SRR1313135 | SRR1313136 | SRR1313137 | SRR1313138 |
| SRR1313139 | SRR1313140 | SRR1313141 | SRR1313142 | SRR1313143 | SRR1313144 |
| SRR1313145 | SRR1313146 | SRR1313147 | SRR1313148 | SRR1313149 | SRR1313161 |
| SRR1313163 | SRR1313164 | SRR1313165 | SRR1313167 | SRR1313170 | SRR1313173 |
| SRR1313174 | SRR1313175 | SRR1313176 | SRR1313178 | SRR1313181 | SRR1313183 |
| SRR1313188 | SRR1313189 | SRR1313190 | SRR1313193 | SRR1313196 | SRR1313197 |
| SRR1313198 | SRR1313201 | SRR1313210 | SRR1313211 | SRR1313212 | SRR1313213 |
| SRR1313214 | SRR1313215 | SRR1313216 | SRR1313217 | SRR1313218 | SRR1313219 |
| SRR1313288 | SRR1313289 | SRR1313290 | SRR1313291 | SRR1313292 | SRR1313293 |
| SRR1313294 | SRR1313295 | SRR1313296 | SRR1313297 | SRR1313300 | SRR1313301 |
| SRR1313302 | SRR1313303 | SRR1373442 | SRR1373444 | SRR1373446 | SRR1373447 |
| SRR1373448 | SRR1373449 | SRR1373457 | SRR1373459 | SRR1374809 | SRR1374811 |
| SRR1374812 | SRR1383358 | SRR1383359 | SRR1383360 | SRR1383361 | SRR1383362 |
| SRR1383363 | SRR1383365 | SRR1383366 | SRR1383367 | SRR1383378 | SRR1383380 |
| SRR1383381 | SRR1383382 | SRR1383383 | SRR1383384 | SRR1383385 | SRR1383386 |
| SRR1383387 | SRR1383388 | SRR1383389 | SRR1383391 | SRR1383392 | SRR1383394 |
| SRR1383395 | SRR1383396 | SRR1383400 | SRR1383401 | SRR1383402 | SRR1383403 |
| SRR1383404 | SRR1383405 | SRR1383406 | SRR1383407 | SRR1383408 | SRR1383409 |
| SRR1409977 | SRR1409978 | SRR1409979 | SRR1474200 | SRR1514140 | SRR1514141 |
| SRR1519064 | SRR1519065 | SRR1519066 | SRR1519067 | SRR1519070 | SRR1525424 |
| SRR1525426 | SRR1525427 | SRR1536577 | SRR1536578 | SRR1538598 | SRR1539402 |
| SRR1539403 | SRR1539405 | SRR1539406 | SRR1544480 | SRR1544481 | SRR1544483 |
| SRR1544488 | SRR1544489 | SRR1544503 | SRR1551084 | SRR1551100 | SRR1551102 |
| SRR1551103 | SRR1551104 | SRR1551105 | SRR1560839 | SRR1560927 | SRR1561865 |

Table 11: SRA Samples (continued)

| | | | | | |
|---|---|---|---|---|---|
| SRR1561866 | SRR1561868 | SRR1561869 | SRR1573501 | SRR1575102 | SRR1575103 |
| SRR1575104 | SRR1575105 | SRR1576140 | SRR1576141 | SRR1576166 | SRR1576168 |
| SRR1576180 | SRR1576181 | SRR1580540 | SRR1580541 | SRR1580542 | SRR1580544 |
| SRR1580545 | SRR1580550 | SRR1580551 | SRR1580552 | SRR1580553 | SRR1582164 |
| SRR1582165 | SRR1582166 | SRR1582167 | SRR1582168 | SRR1582169 | SRR1596208 |
| SRR1596209 | SRR1596210 | SRR1596211 | SRR1596212 | SRR1596213 | SRR1596215 |
| SRR1596216 | SRR1600270 | SRR1600271 | SRR1604987 | SRR1604988 | SRR1604989 |
| SRR1604990 | SRR1604991 | SRR1605004 | SRR1605005 | SRR1605006 | SRR1605007 |
| SRR1609980 | SRR1609981 | SRR1609982 | SRR1609983 | SRR1609984 | SRR1609985 |
| SRR1609986 | SRR1609987 | SRR1609988 | SRR1616899 | SRR1616900 | SRR1616901 |
| SRR1616902 | SRR1616903 | SRR1616904 | SRR1616905 | SRR1616906 | SRR1616907 |
| SRR1616908 | SRR1616909 | SRR1616920 | SRR1616921 | SRR1616922 | SRR1616923 |
| SRR1616924 | SRR1616925 | SRR1616926 | SRR1616927 | SRR1639637 | SRR1639741 |
| SRR1639742 | SRR1639743 | SRR1639744 | SRR1639745 | SRR1643191 | SRR1643192 |
| SRR1643193 | SRR1643194 | SRR1643195 | SRR1643196 | SRR1643197 | SRR1643198 |
| SRR1643199 | SRR1643210 | SRR1643219 | SRR1643230 | SRR1643234 | SRR1643235 |
| SRR1643243 | SRR1643249 | SRR1655002 | SRR1655004 | SRR1655006 | SRR1655010 |
| SRR1657556 | SRR1657557 | SRR1660046 | SRR1660320 | SRR1660321 | SRR1660323 |
| SRR1660324 | SRR1663221 | SRR1663223 | SRR1663224 | SRR1663229 | SRR1663230 |
| SRR1663233 | SRR1663251 | SRR1663253 | SRR1663255 | SRR1663256 | SRR1663257 |
| SRR1663258 | SRR1663259 | SRR1663272 | SRR1663273 | SRR1663275 | SRR1663278 |
| SRR1663288 | SRR1686011 | SRR1691634 | SRR1691648 | SRR1691650 | SRR1691651 |
| SRR1691652 | SRR1691653 | SRR1696812 | SRR1696813 | SRR1696814 | SRR1697330 |
| SRR1697331 | SRR1697334 | SRR1697335 | SRR1697336 | SRR1697337 | SRR1697338 |
| SRR1697339 | SRR1721280 | SRR1721281 | SRR1721282 | SRR1721283 | SRR1721284 |
| SRR1721285 | SRR1721286 | SRR1721301 | SRR1721302 | SRR1721304 | SRR1721305 |
| SRR1721307 | SRR1721308 | SRR1721309 | SRR1737414 | SRR1737419 | SRR1740044 |
| SRR1740068 | SRR1747248 | SRR1747966 | SRR1747967 | SRR1747968 | SRR1747972 |
| SRR1778290 | SRR1778291 | SRR1778293 | SRR1778294 | SRR1778297 | SRR1778299 |
| SRR1778301 | SRR1778302 | SRR1778303 | SRR1778307 | SRR1778308 | SRR1778309 |
| SRR1778310 | SRR1778312 | SRR1778314 | SRR1778315 | SRR1778316 | SRR1781570 |
| SRR1781571 | SRR1803196 | SRR1803197 | SRR1803198 | SRR1803199 | SRR1803200 |
| SRR1803202 | SRR1803203 | SRR1803204 | SRR1803205 | SRR1803206 | SRR1803207 |
| SRR1803211 | SRR1803212 | SRR1803213 | SRR1812361 | SRR1812362 | SRR1812363 |
| SRR1812364 | SRR1812365 | SRR1812366 | SRR1812369 | SRR1946657 | SRR1946659 |
| SRR1946660 | SRR1946662 | SRR1946663 | SRR1946664 | SRR1946665 | SRR1946666 |
| SRR1946667 | SRR1946678 | SRR1946679 | SRR1946680 | SRR1946681 | SRR1946683 |
| SRR1946686 | SRR1946690 | SRR1946691 | SRR1947740 | SRR1949811 | SRR1949813 |
| SRR1949814 | SRR1949816 | SRR1949818 | SRR1949819 | SRR1949830 | SRR1949831 |
| SRR1949832 | SRR1949833 | SRR1949834 | SRR1949835 | SRR1949836 | SRR1949837 |

Table 12: SRA Samples (continued)

| | | | | | |
|---|---|---|---|---|---|
| SRR1949838 | SRR1993909 | SRR1993910 | SRR1993911 | SRR1993912 | SRR1993913 |
| SRR2056365 | SRR2056368 | SRR2071345 | SRR2071347 | SRR2071349 | SRR2071361 |
| SRR2071368 | SRR2071369 | SRR2079880 | SRR2081131 | SRR2081133 | SRR2081137 |
| SRR2081138 | SRR307898 | SRR307904 | SRR307907 | SRR307909 | SRR307910 |
| SRR307911 | SRR307912 | SRR307913 | SRR307915 | SRR307917 | SRR307919 |
| SRR307920 | SRR307921 | SRR307923 | SRR307924 | SRR307925 | SRR307926 |
| SRR307927 | SRR307928 | SRR307929 | SRR307931 | SRR307932 | SRR307933 |
| SRR315113 | SRR315298 | SRR315299 | SRR315300 | SRR315301 | SRR315302 |
| SRR315303 | SRR315305 | SRR315306 | SRR315307 | SRR315308 | SRR315309 |
| SRR315325 | SRR315329 | SRR317035 | SRR317036 | SRR317037 | SRR317038 |
| SRR317039 | SRR317040 | SRR317050 | SRR317051 | SRR317052 | SRR317053 |
| SRR317054 | SRR317055 | SRR317056 | SRR317057 | SRR317058 | SRR317059 |
| SRR317060 | SRR317061 | SRR317062 | SRR317063 | SRR317064 | SRR317065 |
| SRR317066 | SRR317067 | SRR317068 | SRR317069 | SRR332270 | SRR332272 |
| SRR332273 | SRR350717 | SRR350718 | SRR353602 | SRR353603 | SRR364001 |
| SRR364065 | SRR364829 | SRR364830 | SRR364831 | SRR387293 | SRR387393 |
| SRR387395 | SRR387396 | SRR387397 | SRR387398 | SRR387399 | SRR387400 |
| SRR387401 | SRR387402 | SRR387403 | SRR387404 | SRR387405 | SRR387406 |
| SRR387407 | SRR387418 | SRR387426 | SRR387427 | SRR387428 | SRR387429 |
| SRR387430 | SRR387432 | SRR387433 | SRR387434 | SRR387435 | SRR387436 |
| SRR387437 | SRR387438 | SRR387440 | SRR387441 | SRR387442 | SRR387443 |
| SRR387444 | SRR387445 | SRR387446 | SRR387447 | SRR387448 | SRR387521 |
| SRR387661 | SRR387662 | SRR390469 | SRR400342 | SRR400343 | SRR400344 |
| SRR403879 | SRR403880 | SRR403881 | SRR452328 | SRR452329 | SRR452331 |
| SRR452333 | SRR479061 | SRR479070 | SRR486237 | SRR486238 | SRR486239 |
| SRR486240 | SRR486241 | SRR486242 | SRR488136 | SRR488137 | SRR488684 |
| SRR488685 | SRR493937 | SRR493938 | SRR493939 | SRR493940 | SRR493941 |
| SRR493942 | SRR493943 | SRR493944 | SRR493945 | SRR493946 | SRR493947 |
| SRR493948 | SRR493949 | SRR493950 | SRR493957 | SRR493958 | SRR493959 |
| SRR493960 | SRR496581 | SRR496588 | SRR496590 | SRR496593 | SRR496595 |
| SRR496596 | SRR496597 | SRR497884 | SRR497885 | SRR500875 | SRR500876 |
| SRR500877 | SRR500878 | SRR500879 | SRR500880 | SRR500881 | SRR500882 |
| SRR500883 | SRR500884 | SRR500885 | SRR500886 | SRR500887 | SRR500888 |
| SRR500889 | SRR500890 | SRR514854 | SRR514855 | SRR514856 | SRR514857 |
| SRR514858 | SRR514859 | SRR515084 | SRR518000 | SRR521452 | SRR521455 |
| SRR521460 | SRR521462 | SRR521499 | SRR521516 | SRR521521 | SRR521522 |
| SRR521523 | SRR521525 | SRR521526 | SRR521527 | SRR521528 | SRR521529 |
| SRR529646 | SRR529647 | SRR534289 | SRR534290 | SRR534291 | SRR534292 |
| SRR534293 | SRR534294 | SRR534295 | SRR534296 | SRR534297 | SRR534298 |
| SRR534299 | SRR534300 | SRR534301 | SRR534302 | SRR534303 | SRR534304 |

Table 13: SRA Samples (continued)

| | | | | | |
|---|---|---|---|---|---|
| SRR534305 | SRR534306 | SRR534307 | SRR534308 | SRR534309 | SRR534310 |
| SRR534317 | SRR534318 | SRR534323 | SRR534324 | SRR534325 | SRR534326 |
| SRR534327 | SRR534334 | SRR534335 | SRR545685 | SRR545686 | SRR545687 |
| SRR545688 | SRR545689 | SRR545690 | SRR545691 | SRR545692 | SRR545693 |
| SRR545694 | SRR545695 | SRR545696 | SRR545697 | SRR545701 | SRR545702 |
| SRR545704 | SRR545705 | SRR545707 | SRR545708 | SRR545709 | SRR545714 |
| SRR545715 | SRR545716 | SRR545721 | SRR545722 | SRR545723 | SRR545940 |
| SRR545941 | SRR548611 | SRR548612 | SRR548613 | SRR557130 | SRR557131 |
| SRR557133 | SRR557134 | SRR557135 | SRR557136 | SRR557137 | SRR557138 |
| SRR557139 | SRR557140 | SRR567455 | SRR574820 | SRR574821 | SRR574822 |
| SRR574823 | SRR576790 | SRR576791 | SRR576928 | SRR577579 | SRR577580 |
| SRR577581 | SRR577587 | SRR577588 | SRR577589 | SRR577591 | SRR577592 |
| SRR577599 | SRR577600 | SRR577601 | SRR578270 | SRR578627 | SRR578628 |
| SRR578631 | SRR578635 | SRR578640 | SRR578651 | SRR578652 | SRR578755 |
| SRR578756 | SRR585572 | SRR585573 | SRR585574 | SRR585575 | SRR585576 |
| SRR585577 | SRR592574 | SRR592575 | SRR592576 | SRR592577 | SRR592578 |
| SRR592580 | SRR592581 | SRR596103 | SRR597893 | SRR597894 | SRR597895 |
| SRR597900 | SRR597904 | SRR597912 | SRR605543 | SRR606335 | SRR606336 |
| SRR606337 | SRR611842 | SRR611844 | SRR616076 | SRR616077 | SRR616078 |
| SRR616079 | SRR616080 | SRR640257 | SRR640258 | SRR640259 | SRR643741 |
| SRR643742 | SRR643766 | SRR644512 | SRR644513 | SRR644514 | SRR645775 |
| SRR645776 | SRR645777 | SRR645778 | SRR645779 | SRR645780 | SRR645781 |
| SRR645783 | SRR651662 | SRR651663 | SRR651664 | SRR651665 | SRR651666 |
| SRR651669 | SRR651673 | SRR651674 | SRR651675 | SRR651676 | SRR651677 |
| SRR651678 | SRR651687 | SRR651692 | SRR651699 | SRR651700 | SRR651701 |
| SRR651702 | SRR651703 | SRR651704 | SRR651705 | SRR651706 | SRR653217 |
| SRR653218 | SRR653221 | SRR653222 | SRR653391 | SRR653392 | SRR653393 |
| SRR653394 | SRR653395 | SRR653396 | SRR653397 | SRR653398 | SRR653399 |
| SRR653400 | SRR658527 | SRR710084 | SRR710085 | SRR710086 | SRR710087 |
| SRR710088 | SRR710089 | SRR710090 | SRR710091 | SRR710092 | SRR710093 |
| SRR710094 | SRR710095 | SRR748515 | SRR748516 | SRR748517 | SRR748518 |
| SRR748519 | SRR748520 | SRR748521 | SRR748522 | SRR764776 | SRR764777 |
| SRR764778 | SRR764779 | SRR764780 | SRR764781 | SRR764782 | SRR764783 |
| SRR764784 | SRR768411 | SRR768412 | SRR768413 | SRR768414 | SRR770714 |
| SRR770715 | SRR770716 | SRR771465 | SRR771466 | SRR771467 | SRR771547 |
| SRR771548 | SRR771549 | SRR771550 | SRR771553 | SRR771554 | SRR771555 |
| SRR771556 | SRR771557 | SRR771558 | SRR791043 | SRR791044 | SRR791045 |
| SRR791046 | SRR791047 | SRR791048 | SRR791049 | SRR797054 | SRR797055 |
| SRR797056 | SRR797057 | SRR831698 | SRR831699 | SRR831700 | SRR831701 |
| SRR831702 | SRR833716 | SRR833720 | SRR833721 | SRR833722 | SRR833723 |

Table 14: SRA Samples (continued)

| | | | | | |
|---|---|---|---|---|---|
| SRR833724 | SRR833725 | SRR833726 | SRR833727 | SRR833728 | SRR833729 |
| SRR834983 | SRR834985 | SRR835999 | SRR836000 | SRR846952 | SRR921935 |
| SRR921936 | SRR921937 | SRR921939 | SRR921940 | SRR921945 | SRR934360 |
| SRR934361 | SRR934362 | SRR934363 | SRR934364 | SRR934365 | SRR934366 |
| SRR934367 | SRR934368 | SRR934369 | SRR934641 | SRR934643 | SRR934645 |
| SRR934717 | SRR934720 | SRR934730 | SRR934731 | SRR934732 | SRR934733 |
| SRR934734 | SRR934735 | SRR934740 | SRR934741 | SRR934742 | SRR934746 |
| SRR934750 | SRR934751 | SRR934753 | SRR934766 | SRR934769 | SRR934770 |
| SRR934771 | SRR934772 | SRR934778 | SRR934781 | SRR934782 | SRR934784 |
| SRR934785 | SRR934787 | SRR934790 | SRR934793 | SRR934794 | SRR934797 |
| SRR934800 | SRR934801 | SRR934802 | SRR934804 | SRR934808 | SRR934809 |
| SRR934810 | SRR934815 | SRR934817 | SRR934826 | SRR934827 | SRR934829 |
| SRR934836 | SRR934839 | SRR934844 | SRR934846 | SRR934849 | SRR934855 |
| SRR934856 | SRR934859 | SRR934867 | SRR934873 | SRR934875 | SRR934879 |
| SRR934881 | SRR934882 | SRR934883 | SRR934887 | SRR934889 | SRR934890 |
| SRR934894 | SRR934895 | SRR934906 | SRR934907 | SRR934908 | SRR934910 |
| SRR934911 | SRR934914 | SRR934915 | SRR934916 | SRR934926 | SRR934927 |
| SRR934928 | SRR934930 | SRR934931 | SRR934933 | SRR934934 | SRR934935 |
| SRR934937 | SRR934943 | SRR934945 | SRR934947 | SRR934949 | SRR934954 |
| SRR934958 | SRR934959 | SRR934964 | SRR934966 | SRR934968 | SRR934970 |
| SRR934971 | SRR934972 | SRR934974 | SRR934977 | SRR934978 | SRR934979 |
| SRR934990 | SRR935375 | SRR935376 | SRR935670 | SRR935672 | SRR935673 |
| SRR935674 | SRR935675 | SRR935676 | SRR935677 | SRR935678 | SRR935679 |
| SRR935680 | SRR935681 | SRR941857 | SRR941858 | SRR941859 | SRR941860 |
| SRR941861 | SRR950079 | SRR962599 | SRR962602 | SRR972707 | SRR972709 |
| SRR980480 | SRR988501 | SRR988507 | SRR988509 | SRR988510 | |
